# Learning dynamics by computational integration of single cell genomic and lineage information

**DOI:** 10.1101/2021.05.06.443026

**Authors:** Shou-Wen Wang, Allon M. Klein

## Abstract

A goal of single cell genome-wide profiling is to reconstruct dynamic transitions during cell differentiation, disease onset, and drug response. Single cell assays have recently been integrated with lineage tracing, a set of methods that identify cells of common ancestry to establish *bona fide* dynamic relationships between cell states. These integrated methods have revealed unappreciated cell dynamics, but their analysis faces recurrent challenges arising from noisy, dispersed lineage data. Here, we develop coherent, sparse optimization (CoSpar) as a robust computational approach to infer cell dynamics from single-cell genomics integrated with lineage tracing. CoSpar is robust to severe down-sampling and dispersion of lineage data, which enables simpler, lower-cost experimental designs and requires less calibration. In datasets representing hematopoiesis, reprogramming, and directed differentiation, CoSpar identifies fate biases not previously detected, predicting transcription factors and receptors implicated in fate choice. Documentation and detailed examples for common experimental designs are available at https://cospar.readthedocs.io/.

## Introduction

In tissue development, regeneration, and disease, cells differentiate into distinct, reproducible phenotypes. A ubiquitous challenge in studying these processes is to order events occurring during differentiation^1–3^, and to identify events that drive cells towards one phenotype or another. This challenge is common to understanding mechanisms in embryo development, stem cell self-renewal, cancer cell drug resistance, and tissue metaplasia^1–3^.

At least two observational strategies help to order cellular events. Single-cell genome-wide profiling – such as by single-cell RNA sequencing (scRNA-seq) – offers a universal and scalable approach to observing dynamic states by densely sampling cells at different stages^3–10^. However, scRNA-seq alone does not identify which early differences between cells drive or correlate with fate^2,11–13^. Conversely, lineage tracing offers a complementary family of methods that can clarify long-term dynamic relationships across multiple cell cycles. To carry out lineage tracing, individual cells are labeled at an early time point^1–3^. The state of their clonal progeny is analyzed at one or more later time points (Fig. 1**a**).

**Fig. 1.**
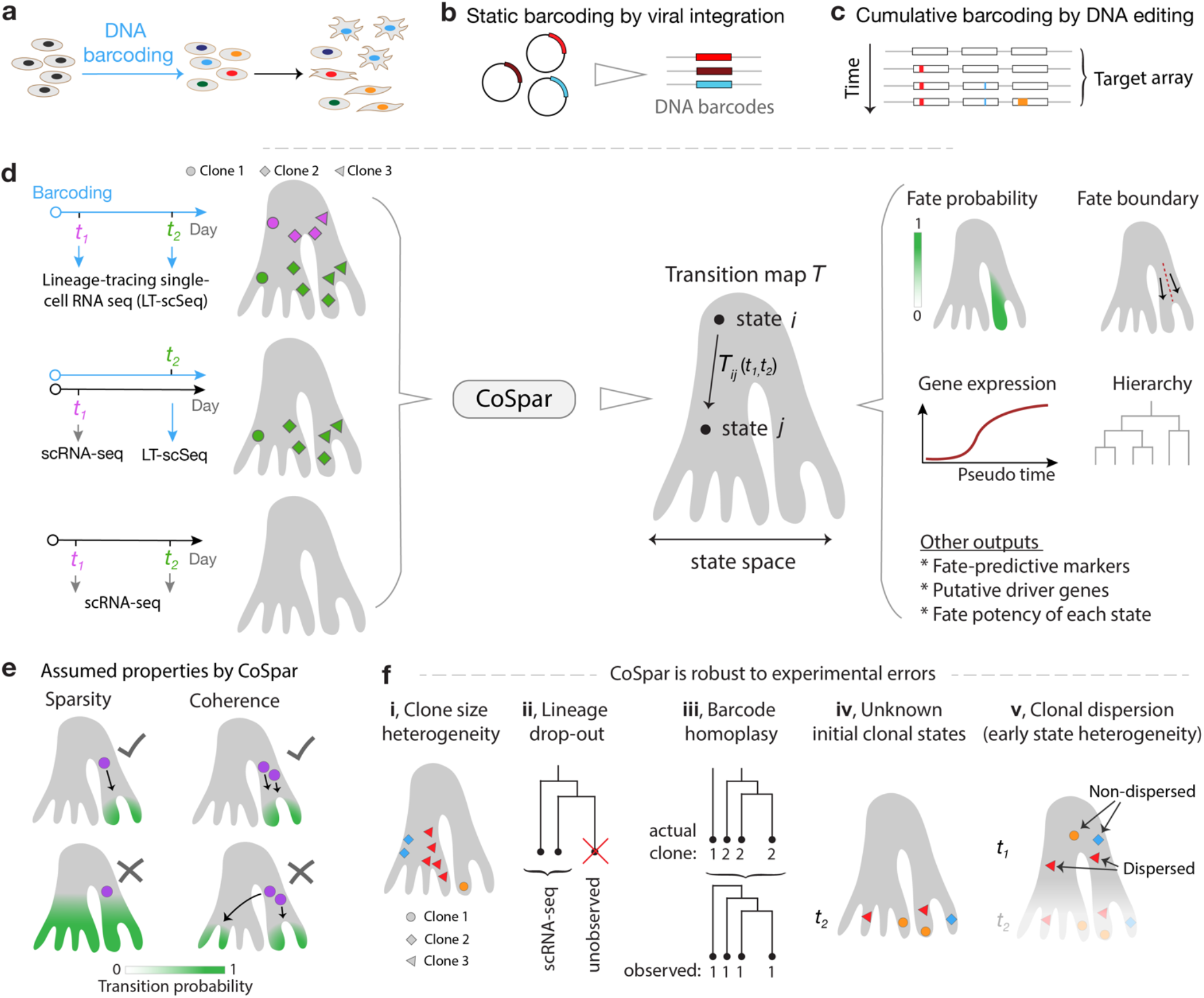
Integrative analysis of lineage tracing and transcriptome data. **a**, Lineage-tracing single cell genomics (LT-scSeq) experiments simultaneously measure cell phenotypes and clonal lineage (indicated by colors). **b-c**, LT-scSeq assays encode lineage information with static DNA barcodes or cumulative barcoding. **d**, CoSpar unifies analysis of different experimental designs to infer transition maps (see text) to reveal fate boundaries, lineage hierarchy, putative markers, and putative fate-determinants. Here and below, the shaded gray regions schematically show a manifold of observed single cell genomic states. **e**, Two key assumptions constrain dynamic inference by CoSpar. **f**, Stereotypical challenges in clonal analysis. Single labeled cells can give rise to clones with a wide dispersion in size; LT-scSeq loses cells during analysis leading to loss of clonal structure; barcode homoplasy occurs when cells from different clones present the same barcode due to experimental limitations; progenitor states are not observed when clones are only observed upon tissue dissociation; clonal dispersion occurs when early clonal states are heterogeneous due to the lag time between barcoding and profiling.

Recently, a number of efforts from us and others have integrated lineage-tracing with single-cell genome-wide profiling (hereafter LT-scSeq) using unique, heritable, and expressed DNA barcodes^2,13–21^. These technologies identify cells that share a common ancestor and define their genomic state in an unbiased manner. LT-scSeq experiments have been used to successfully identify when fate decisions occur^13,14^, novel markers for stem cells^16^, and pathways which control cell fate choice^14,16^. The simplest of these methods labels cells at one time point^13^ (Fig. 1**b**); more complex methods allow the accumulation of barcodes over successive cell divisions to reveal the substructure of clones^2,13–21^ (Fig. 1**c**).

Emerging LT-scSeq methods have been successful at revealing novel regulators of cell fate^14,16^ and the fate potential of early progenitors^13,14^, but they also present challenges that may limit their utility in practice. We identified at least five technical and biological challenges that affect experimental design and interpretation (Fig. 1**f**). These include stochastic differentiation and variable expansion of clones^22^ (Fig. 1**f-i**), cell loss during analysis (Fig. 1**f-ii**), barcode homoplasy wherein cells acquire the same barcode despite not having a lineage relationship^2^ (Fig. 1**f-iii**), access to clones only at a single time point^23,24^ (Fig. 1**f-iv**), and clonal dispersion due to a lag time between labeling cells and the first sampling (Fig. 1**f-v**). Addressing these problems should greatly simplify the design and interpretation of LT-scSeq assays and put them in the hands of a wider research community. To our knowledge, there is not yet an analysis method that systematically overcomes these problems.

Here, we develop a robust and generalizable computational approach to analyze LT-scSeq experiments. We begin with a model of clonal dynamics in which cells divide, differentiate, or are lost from the sampled tissue in a stochastic manner, with rates that are state-dependent (Supplementary Fig. 1**a**). We use this model to learn from the data the fraction of progeny of cells, initially in one state, which are found to occupy a second state after some time interval (Fig. 1**d**, Supplementary Fig. 1**b,c**). Our approach captures differentiation bias and fate hierarchies, and can reveal genes whose early expression is predictive of future fate choice.

## Results

### Dynamic inference from clonal data with state information

A formalization of dynamic inference is to identify a transition map, a matrix *T*_*ij*_ (*t*_1_, *t*_2_)^7,25^. We define *T*_*ij*_ (*t*_1_, *t*_2_) specifically as the fraction of progeny of a cell, initially in some state *i* at time *t*_1_, that occupy state *j* at time *t*_2_ (Fig. 1**d**, Supplementary Fig. **1c**). This transition matrix represents a coarse-grained view of the cell dynamics : it already combines the effects of cell division, loss, and differentiation (Supplementary Fig. **1d**). As will be seen, even learning *T*_*ij*_ (*t*_1_, *t*_2_) will prove useful for several applications (Fig. 1**d**).

We make two reasonable assumptions about the nature of biological dynamics to constrain inference of the transition map. We assume the map to be a sparse matrix, since most cells can access just a few states during an experiment (Fig. 1**e**, left panel). And we assume the map to be locally coherent, meaning that cells in similar states should share similar fate outcomes (Fig. 1**e**, right panel). These constraints together force transition maps to be parsimonious and smooth, which makes them robust to practical sources of noise in LT-scSeq experiments (Supplementary Fig. 1**e**). Box 1 formalizes the two constraints and lays out the technical foundation for inferring a transition map by coherent sparse (CoSpar) optimization (see schema in Fig. 2**a**; Supplementary Fig. 2). As inputs, CoSpar requires a clone-by-cell matrix *I*(*t*) that encodes the clonal information at time *t*, and a data matrix for observed cell states (e.g. from scRNA-seq).

**Fig. 2.**
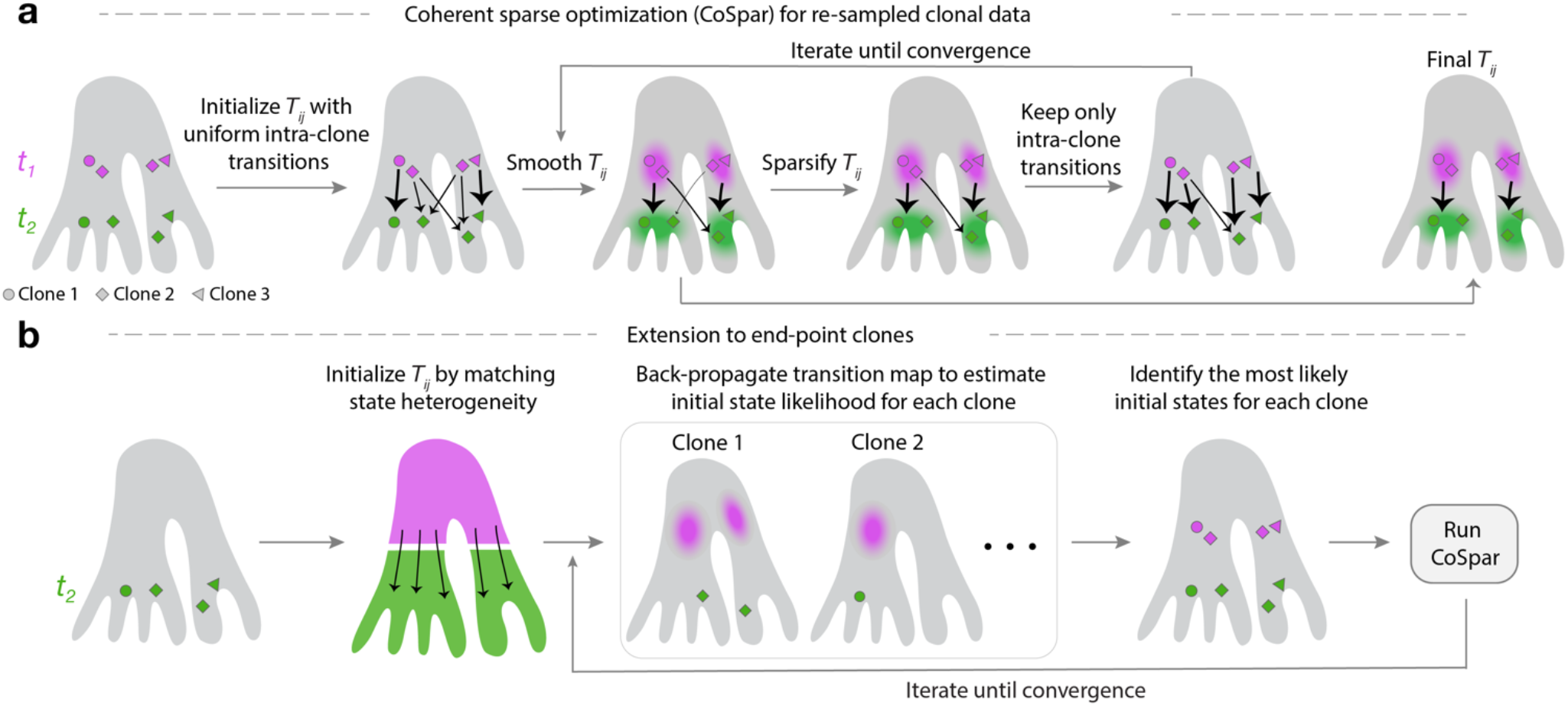
The CoSpar algorithm. **a**, When clones are resampled at two time points, a transition map is inferred by iteratively enforcing observed clonal transitions, coherence (by smoothing) and sparsity until convergence is achieved. (See details and derivation in Methods and Supplemental Note 3). **b**, When clones are observed only once, we infer their progenitor fate bias and identity by first initializing a transition map without clonal information, then iteratively (1) back-propagating the map to predict clonal progenitor identity and (2) learning the transition map as in **a** Until the map and progenitor identities jointly converge.

CoSpar is formulated assuming that we have information on the same clones at more than one time point. More often, one might observe clones at only one time point *t*_2_. For these cases CoSpar jointly optimizes the transition map *T* and the initial clonal data *I*(*t*_1_) (Fig. 2**b**; Methods). In this joint optimization, one must initialize the transition map; we have shown that the final result is robust to initialization (Supplementary Fig. 3**e**; Supplementary Fig. 4**c**,**d**). This approach can be used for clones with nested structure (Supplementary Fig. 4**f**-**h**). Finally, coherence and sparsity provide reasonable constraints to the common problem of predicting dynamics from state heterogeneity alone without lineage data^7^. We extended CoSpar to this case. Thus, CoSpar is flexible to different experimental designs, as summarized in Fig. 1**d**.

#### Box 1

##### Coherent Sparse Optimization

In a model of stochastic differentiation, cells in a clone are distributed across states with a time-dependent density profile *P*(*t*). A transition map *T* directly links clonal density profiles *P* (*t*_1,2_) between time points:

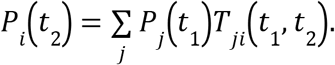

From multiple clonal observations, our goal is to learn *T*. To do so, we denote *I*(*t*) as a clone-by-cell matrix and introduce *S* as a matrix of cell-cell similarity over all observed cell states, including those lacking clonal information. The density profiles of all observed clones are estimated as *P*(*t*) ≈ *I*(*t*)*S*(*t*).

With enough clonal information, *T*(*t*_1_, *t*_2_) could in principle be learnt by matrix inversion. However, the number of clones will always be far less than the number of states. To constrain the map, we require that: 1) *T* is a sparse matrix (Fig. 1**e**, left panel); 2) *T* is locally coherent (Fig. 1**e**, right panel); and 3) *T* is a non-negative matrix. With these requirements, the inference can be formulated as the following optimization problem:

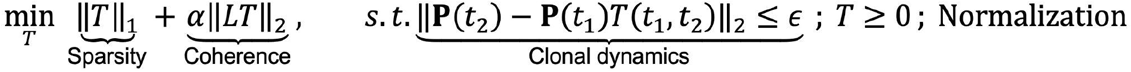

‖*T*‖_1_ quantifies the sparsity of the matrix *T* through its L1 norm, and ‖*LT*‖_2_ quantifies the local coherence of *T* (*L* being the Graph Laplacian of the cell state similarity graph, and *LT* being the local divergence). The remaining constraints enforce the observed clonal dynamics, non-negativity of *T*, and map normalization, respectively. At α = 0, the minimization takes the form of *Lasso*^26^, an algorithm for compressed sensing. Our formulation extends compressed sensing from vectors to matrices, and to enforce local coherence. The local coherence extension is reminiscent of the *fused Lasso* problem^27^. An iterative, heuristic approach solves the CoSpar optimization efficiently (Fig. 2**a**; Supplementary Fig. 2). See Methods and Supplementary Notes 1-3 for further details.

Computer simulations validate that CoSpar recovers dynamics with quantitative accuracy, and they establish that CoSpar inference is robust to two errors typical of LT-scSeq -- barcode homoplasy and clonal dispersion. We modeled cells progressing through a sequence of gene expression states either towards a single fate (Fig. 3**a**) or bifurcating into two fates (Fig. 3**e**), with clones sampled in a manner representative of LT-scSeq experiments^13,14^. With 1000 clones – typical of real experiments – mean transition rates inferred by CoSpar were within 3 standard deviations of the actual transition rate 98% of the time (TPR>98%, Fig. 3**d**) and the distribution of progeny fates showed 85% Pearson correlation to ground truth (Fig. 3**j**). Inferences remained similarly accurate with as few as 30 clones (Fig. 3**d**). CoSpar was robust to barcode homoplasy, and only detectably lost accuracy when all lineage barcodes mixed more than ten clones on average (Fig. 3**a-d**). This degree of homoplasy is far higher than expected in most experiments. Further, CoSpar was robust to clonal dispersion, simulated by sampling clones at increasing times post-barcoding (Fig. 3**f-i**). Conversely, approaches used in previous work, which average the transitions between cells observed in each clone at different time points^13^, are severely affected by both lag time and barcode homoplasy (Fig. 3**d,g,i**).

**Fig. 3.**
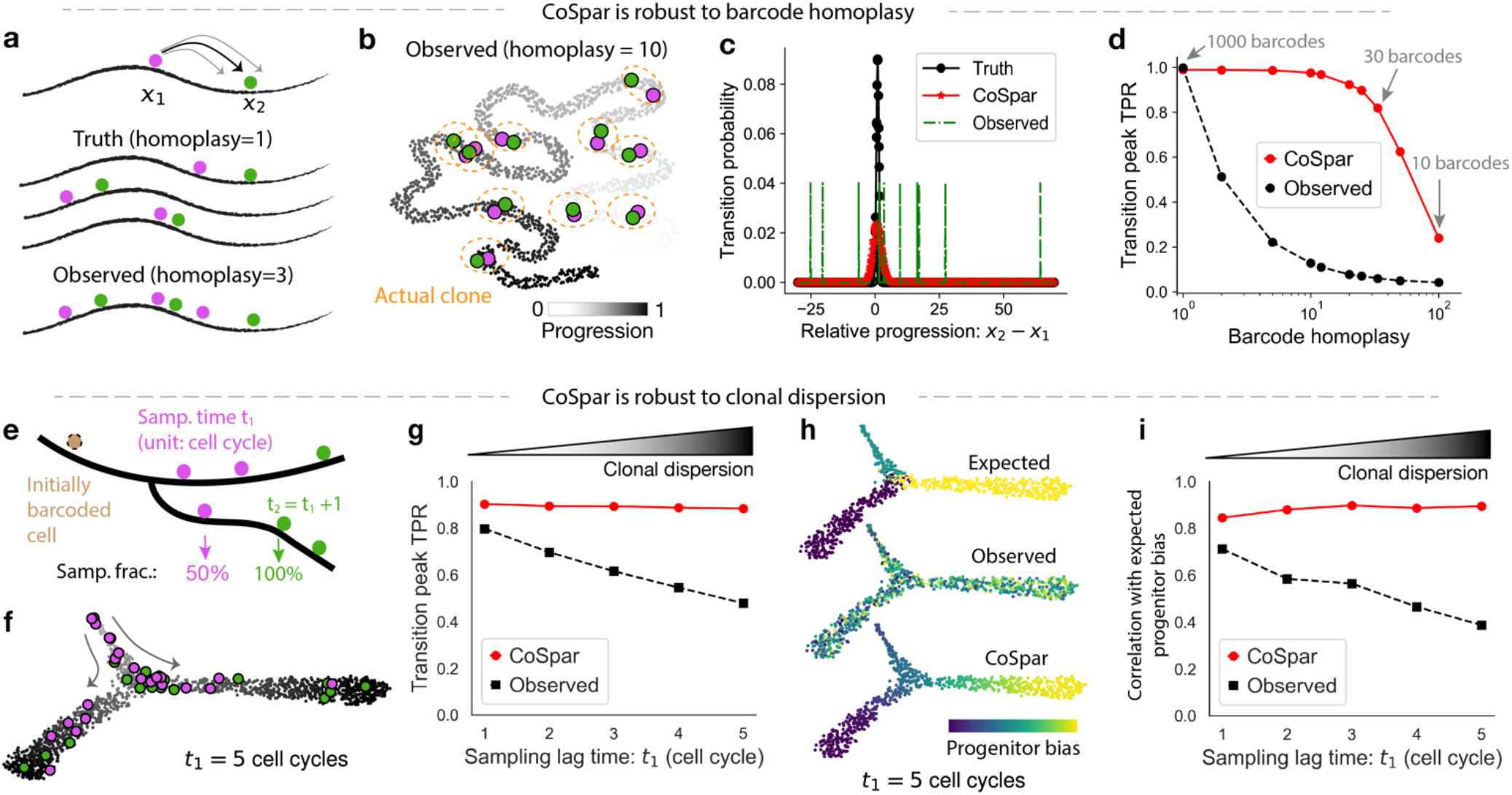
Proof-of-concept with simulated data. **a-d**, Benchmarking transition map inference with barcode homoplasy errors. **a**, Schematics of a simplified simulated LT-scSeq experiment to evaluate the accuracy of CoSpar and its robustness to barcode homoplasy errors. Homoplasy is simulated by assigning multiple clones with the same barcode. **b**, UMAP embedding of simulated data. Cells labeled with one barcode are shown, with moderate homoplasy (10 clones / barcode). **c**, Distribution of true and inferred transition map matrix elements. Observed transitions are broadly distributed due to homoplasy errors, which associate progenitor cells and their progeny across different clones. CoSpar suppresses such transitions by enforcing sparsity and coherence. **d**, CoSpar is robust to severe barcode homoplasy, as seen from the fraction of predicted transitions within 3 standard deviations of the true peak (TPR). **e-i**, Benchmarking transition map inference with clonal dispersion. **e**, Schematics of a second simulated LT-scSeq experiment including variable lag times between clonal labeling and observation. **f**, UMAP embedding of simulated data, with one example clone shown. The clone is first observed 5 cell divisions after initial labeling. **g**, Quantitative evaluation of dynamic inference as a function of the sampling lag time. Growing lag time leads to higher clonal dispersion. Legend and transition peak TPR are defined as in **d**. **h**, Progenitor bias evaluated from the true and inferred transition maps with a simulated sampling lag time of five cell cycles. All clones are highly dispersed, providing no observed bias among early and late states; imposing sparsity enables recovering the true bias. **i**, Quantification of the correlation between true and inferred progenitor bias (shown in **h**), over different sampling lag times.

### CoSpar predicts early fate bias in hematopoiesis

We applied CoSpar to published datasets from three independent experiments. The first experiment tracked hematopoietic progenitor cells (HPCs) differentiating in culture, with clones sampled on days 2, 4 and 6 post-barcoding (Fig. 4**a,b**)^13^. During this time, cells progressed from a heterogeneous pool of HPC states into ten identifiable differentiated cell types. We used all clonal data to generate a ground truth for the early fate bias towards either the monocyte or neutrophil fate, using the method from Weinreb et.al.^13^ (Fig. 4**c**).

**Fig. 4.**
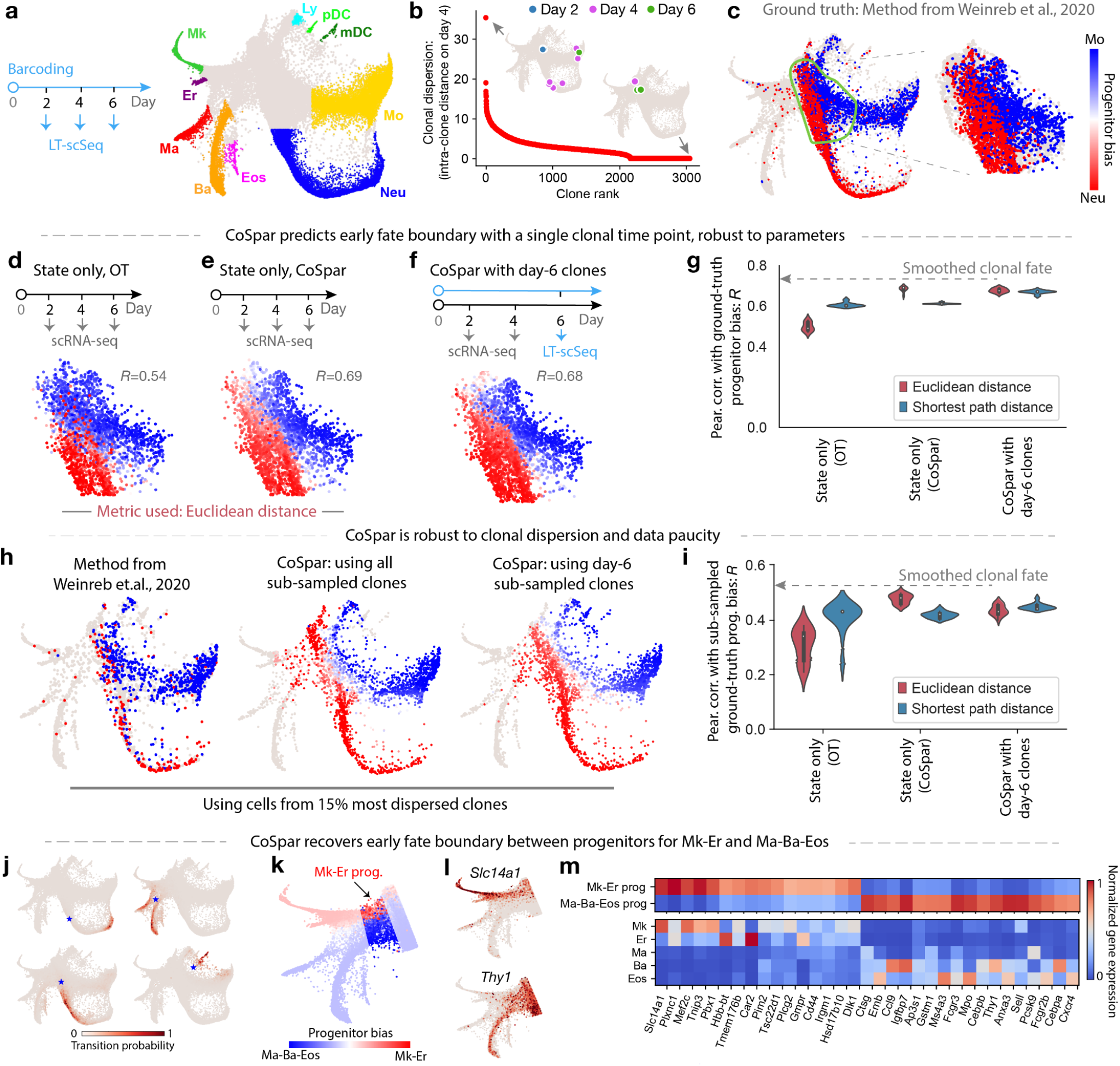
Benchmarking CoSpar and prediction of progenitor bias in hematopoiesis. **a**, Experimental design and SPRING visualization of the hematopoiesis dataset from Weinreb et al.^13^. Early hematopoietic progenitors differentiate into megakaryocyte (Mk), erythrocyte (Er), mast cell (Ma), basophil (Ba), eosinophil (Eos), neutrophil (Neu), monocyte (Mo), lymphoid precursor (Ly), migratory (ccr7+) dendritic cells (mDC), plasmacytoid DC (pDC). **b**, Clones ranked by intra-clone dispersion (i.e., mean intra-clone graph distance) over the observed cell states after 4 days of differentiation. Two illustrative clones are shown. **c**, Bias towards Mo or Neu fate evaluated from all clonal data using the original method in Weinreb et al^13^. Bias among early progenitors (right panel) serves as ground truth for benchmarking. **d,e**, Baseline inference of progenitor bias using optimal transport (OT) or CoSpar, using only state information but no clonal data. **f**, CoSpar inference of progenitor bias using clonal data from a single-clonal time point. **g**, Violin plot showing the distribution of fate prediction outcomes, quantified by the Pearson correlation of the inferred fate bias with the ground truth. The distribution reflects differences in parameters for the OT method (which is used to initialize CoSpar) and choice of distance metric used, showing that clonal data reduces sensitivity to parameter choices in data analysis. Dashed line shows the upper limit expected from cross-validation of benchmarking. **h**, Fate bias inferred using only the 15% most dispersed clones (ranked in panel **b**). **i**, Violin plots showing the distribution in inference performance with the down-sampled data (quantified as in **f**) across parameter values. **j-m**, Predicting the transcriptional identity of Gata1^+^ Mk-Er and Ma-Ba-Eos progenitors using CoSpar. **j**, Representative values of the inferred transition map for 2-day transitions from 4 example cell states (indicated by *). **k**, Heat map of predicted progenitor bias towards Mk-Er and Ma-Ba-Eos fates, overlaid on the state embedding. **l**, Expression of selected genes correlating strongly with predicted fate bias. **m**, Expression heat map for selected genes differentially expressed between the Mk-Er and Ma-Ba-Eos progenitors. Full list of fate-associated genes is provided in Supplementary Table 1.

As a baseline for comparison, we applied CoSpar to predict HPC fate bias using state information alone (Fig. 4**e**). For this and further comparisons, we report the accuracy of fate prediction using Pearson correlation of predicted fate bias with that observed using all clonal data (‘ground truth’). Even without access to any clonal data, CoSpar could resolve early fate bias at a performance close to the upper bound defined by cross-validation of the ground-truth data (CoSpar correlation R=0.69; ground-truth R=0.72) (Fig. 4**e,g**; Supplementary Fig. 6**a**). This performance reflects improvements from enforcing coherence and sparsity (R=0.51-0.54 prior to CoSpar; Fig. 4**d**; Supplementary Fig. 3**f**). However, the prediction based on state information alone is limited because it is sensitive to the choice of distance metric used in analysis (Fig. 4**g**; Supplementary Fig. 3**e**).

Clonal information eliminated the sensitivity to distance metric. To show this, we applied CoSpar to data restricted in time, or restricted in its quality and depth. Using even a single time point of clonal data (day 6), CoSpar recovered early fate bias (Fig. 4**f**; R=0.68), and it did so robustly over a range of parameters and choices of distance metrics (Fig. 4**g**; Supplementary Fig. 3**e**). Further, it recovered the differentiation hierarchy seen in the correlation of clonal barcodes across all cell types (Supplementary Fig. 3**c,d**). When using a sub-sampled dataset from the top 15% most dispersed clones as ranked by day 4 intra-clone distance (Fig. 4**b**), CoSpar performed similarly well, and outperformed the method from Weinreb et al., which was used to analyze this data originally^13^ (Fig. 4**h,i**; Supplementary Fig. 3**a,b**). Thus, CoSpar successfully facilitates analysis of clones at a single time point, or using a fraction of the original data collected in this example.

These benchmarks suggest that CoSpar should be able to predict fate biases not previously recognized. We investigated fate biases in the *Gata1*^+^ states that give rise to five mature fates: megakaryocyte (Mk), erythrocyte (Er), mast cell (Ma), basophil (Ba), and eosinophil (Eos) (Fig. 4**a,k**). In culture, Mk and Er arise from a common progenitor (MEP), and Ba, Eos and Ma are produced by a different progenitor (BEMP)^30,31^. Existing studies of these progenitors are hampered by the lack of good markers. While molecular signatures of FACS-sorted MEP have been explored recently^32^, less is known about the transcriptomic identity of BEMPs. This dataset provides an opportunity to predict the molecular identity of these early progenitors. The original method used to analyze this data finds very few genes distinguishing BEMPs and MEPs (Supplementary Fig. 3**g-i**). Applying CoSpar, we predict an early fate decision boundary between MEP and BEMPs (Fig. 4**j,k**), which correlates with the early expression of genes later associated with the resulting cell types (*Slc14a1* for Mk^32^, *Thy1* for Ba^33^; Fig. 4**l**), and with the transcription factor (TF) *Cebpa* that regulates Eos and Ba differentiation^30^. We identified 377 known and novel putative fate-associated genes (Fig. 4**m**; Supplementary Table 1). Differences between the putative BEMPs and MEPs are evident in scRNA-seq data, and clonal data integrated by CoSpar supports that the differences are associated with functional fate bias. This analysis highlights that CoSpar can identify fate-predictive genes from limited LT-scSeq data.

### CoSpar reveals early fate bias in reprogramming

The second experiment we analyzed tracked cells during the reprogramming of fibroblast cells over 28 days into endodermal progenitors (Fig. 5**a**)^14^. In this experiment, approximately 30% of cells successfully reprogrammed; the remainder failed. Clonal analysis with cumulative barcoding was used to identify these cells early and predicted features that regulate their fate (Fig. 5**b,c**). We used clones strongly enriched in one of the two fates, identified by the original study, to generate the ground truth for early fate bias, and we then used it to benchmark CoSpar.

**Fig. 5.**
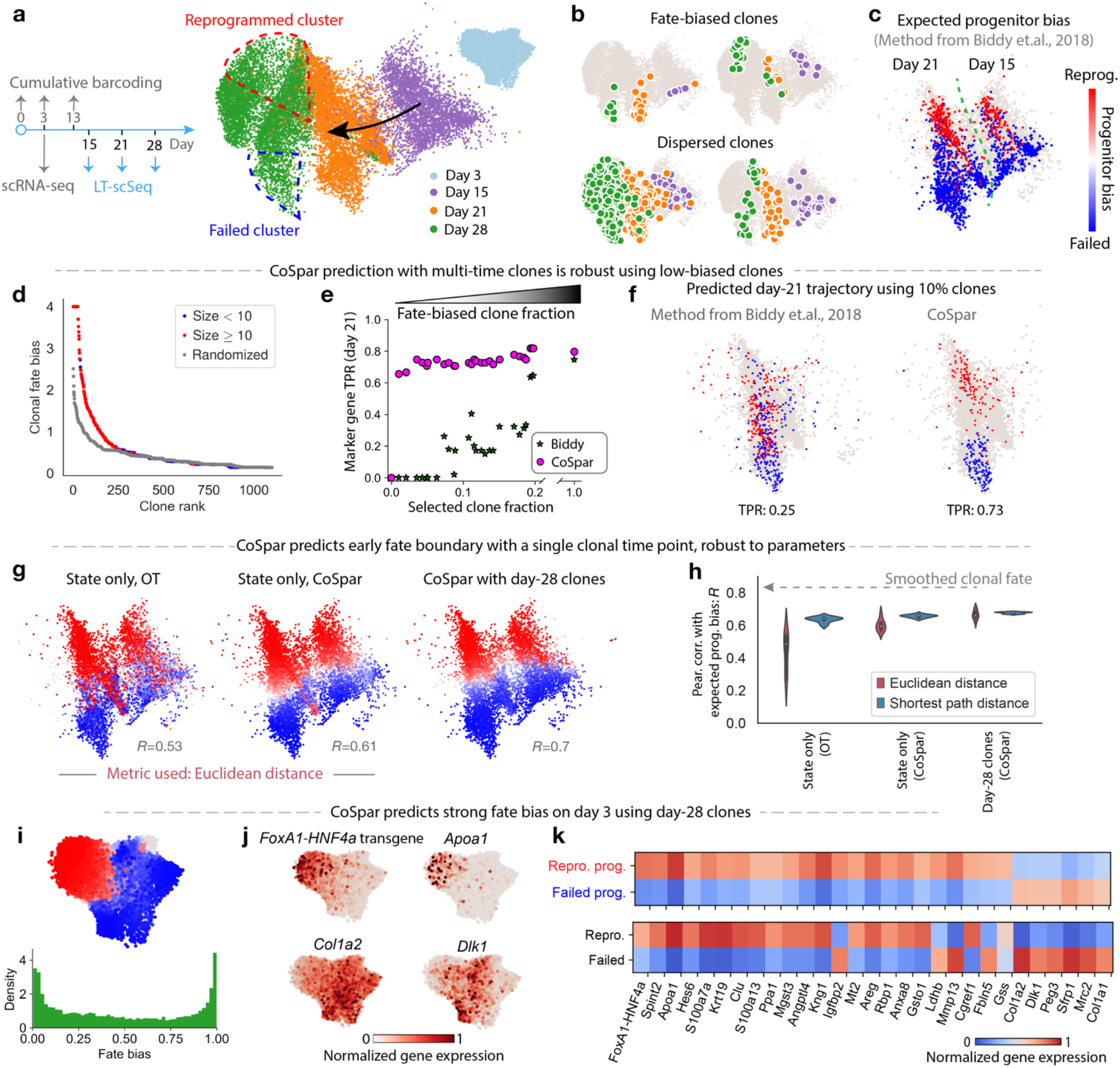
Progenitor bias in fibroblast reprogramming. **a**, Experimental design and UMAP visualization of cell reprogramming from fibroblast cells to induced endoderm progenitors (iEP) by ectopic expression of a transgene *FoxA1-HNF4a* on day 0^14^. Schema shows time points for scRNA-seq only (grey arrows) and LT-scSeq (blue arrows). **b**, The UMAP visualization overlaid with examples of individual clones. Cells are colored by time point as in **a**. **c**, UMAP visualization of transcriptomes on days 15 and 21 of reprogramming, colored by progenitor bias towards successful or failed reprogramming fates, using cells in clones selectively filtered for strong fate bias as in the original study^14^. **d-f**, Benchmarking CoSpar using clones with weak fate bias. **d**, Clones ranked by consistency in the fate outcomes of their constituent cells [fate bias defined as −log(p-value), Fisher Exact test]. **e**, Accuracy in predicting the fate outcome of cells observed on day 21 using data from progressively fewer fate-biased clones. Predictions use the original method (Biddy et al.^14^) or CoSpar. Accuracy is assessed in the true positive rate (TPR) of identifying genes associated with fate outcomes previously reported in Ref^14^. **f**, UMAP visualization showing the cell states on day 21 predicted to undergo successful or failed reprogramming, when using the 10% clones with lowest fate bias. **g-h**, CoSpar predicts early progenitor bias with a single clonal time point, robust to parameters. **g**, Progenitor bias on days 15 and 21 predicted using only state information; or with end-point (day 28) clonal information only. **h**, Violin plots as in Fig. 4**g** Quantifying prediction accuracy over a range of parameters, showing consistent improvement by imposing coherence, sparsity, and enforcing clonal relationships. **i-k**, Predicting early fate determination within 3 days of transgene expression. **i**, Predicted progenitor bias of cells on day 3. **j**, Expression on day-3 states of selected genes predicted to correlate with successful or failed reprogramming. **k**, Expression of additional genes differentially expressed on day 3 between cells predicted to succeed or to fail reprogramming. See the full list at Supplementary Table 2.

To evaluate CoSpar, we revisited this experiment after discarding over 90% of clones, and we specifically retained clones that show the least bias in reprogramming outcomes. Despite deliberately using down-sampled low-quality data, CoSpar recapitulated fate bias: the predicted progenitors of reprogrammed and failed cells share 73 out of 100 marker genes with the ground truth population (Fig. 5**f**), including genes previously showing strong positive and negative association with reprogramming success (*Apoa1*, *Spint2*, *Col1a2*, *Peg3*), as well as *Mettl7a1*, which was found to improve reprogramming^14^. These genes could be associated with fate bias using as few as ten clones, even when deliberately selecting clones with minimal fate bias (Fig. 5**d,e**; Supplementary Fig. 4**b**). By contrast, the analytical approach used in the original study^14^ failed to identify fate-predictive gene expression after such severe reduction in data quality (Fig. 5**e,f**; Supplementary Fig. 4**b**). Further, CoSpar performed robustly when using only clonal data from the final time point of the experiment (Fig. 5**g,h**; Supplementary Fig. 4**c-e**).

As in hematopoiesis, it is instructive to see the information encoded in clonal relationships. When applying CoSpar without clonal data, we found that CoSpar could predict the same early fate biases (Fig. 5**g**, Middle panel), but is again sensitive to the distance metric used (Fig. 5**h**). A different distance metric performs best here from the hematopoiesis dataset, suggesting that there is no simple ‘best-practice’ approach to dynamic inference in the absence of clonal data.

Finally, we applied CoSpar to predict fate bias at the earliest available time point after reprogramming is initiated (day 3), where no clonal information is available and fate bias remains unexplored^14^. Using clonal information, CoSpar predicts strong fate biases (Fig. 5**i**), arguing that future reprogramming success is established very early on. This prediction is supported by the differential expression of transgene *FoxA1-HNF4a* (a TF cocktail to induce reprogramming), the reprogramming marker gene *Apoa1*, and failed trajectory marker *Col1a2* and *Dlk1*^14^ (Fig. 5**j)**. We also identified multiple genes predicted to correlate with fate bias on day 3 and whose significance in reprogramming has not been previously established (Fig. 5**k**; Supplementary Table 2).

### CoSpar predicts early fate bias during lung directed differentiation

In the third experiment, human pluripotent stem cells were differentiated into distal lung alveolar epithelial cells (induced alveolar epithelial type 2 cells, or iAEC2s)^23,34^. Here, clonal and transcriptomic information were profiled jointly on day 27 after initial barcoding on day 17, and a separate time-course experiment produced scRNA-seq data for 6 time points, including days 17 and 21(Fig. 6**a**). In this study, Hurley et al. reported the existence of clones derived from multipotent cells on day 17 but did not investigate their fate biases. A re-examination of the clonal data, however, suggests strong fate biases as early as day 17. Out of the 272 clones, 25% were enriched in either the iAEC2 or non-iAEC2 clusters (FDR=0.01), and clonal compositions differed significantly from that of randomized clones (Fig. 6**b**). Accordingly, clonal representation of iAEC2s anti-correlates with other fates (Supplementary Fig. 5**b,c**). We investigated signatures that could predict effectors of fate bias among day 17 progenitors.

**Fig. 6.**
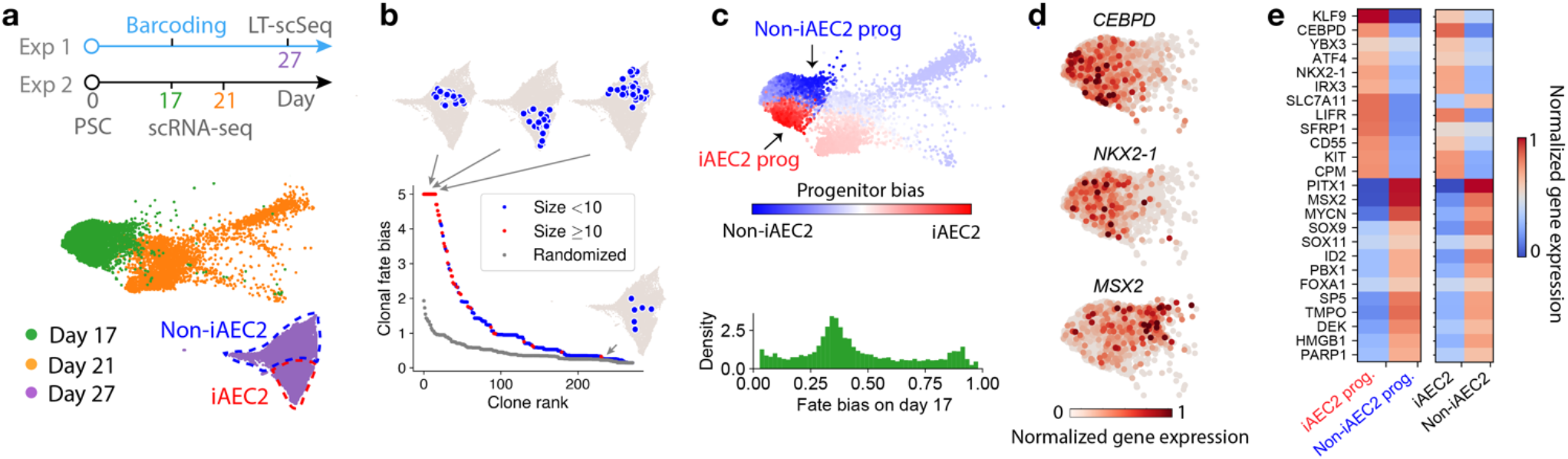
Progenitor bias during hPSCs differentiation into endodermal lineages. **a**, Experimental design and UMAP visualization for differentiating human pluripotent stem cells (hPSC) into induced alveolar epithelium (iAEC2) lung cells and other endodermal cell types. **b**, Clones ranked by fate bias towards iAEC2 fate (bias defined as in Fig. 5**d**), with representative biased (top) and dispersed (bottom) clones shown. **c**, Predicted progenitor bias of cells towards iAEC2 fate on day 17 of differentiation, overlaid on the state embedding and shown as a histogram. **d**,**e** Expression on day-17 states of selected genes predicted to correlate with iAEC2 and non-iAEC2 fates. In **e**, expression is shown alongside the corresponding expression in mature cells on day 27.

Applying CoSpar, we assigned a putative fate bias to each of the cells seen on day 17. CoSpar predicts some cells to be strongly biased in cell fate (Fig. 6**c**), and also the existence of unbiased multipotent states; these strongly overlap with highly proliferating cell states on day 17 and are consistent with large clones hosting multiple endodermal lineages on day 27 (Supplementary Fig. 5**d**). As a control, we expected weaker fate biases earlier in differentiation, which is confirmed by applying CoSpar to cells two days earlier (day 15, Supplementary Fig. 5**e-g**). Among genes differentially expressed between the two biased populations on day 17, we identified several established TFs that regulate lung differentiation: *CEBPD, NKX2-1, SOX9, SOX11* (Fig. 6**d,e**; Supplementary Table 3)^23,35–37^.

## Discussion

Here we have developed a computational framework for systematically inferring dynamic transitions by integrating state and clonal information. It extends the problem of compressed sensing. Our method takes advantage of reasonable assumptions on the nature of biological dynamics: that cells in similar states behave comparably, and that cells limit their possible dynamics to give sparse transitions. Using published datasets, we demonstrated that coherent sparse optimization relates molecular heterogeneity of cells to their future fate outcomes in a manner that is robust to typical sources of experimental error (Fig. 1**f**), using as little as 5-10% data originally collected in prior experiments. The computational methods used in each original study to analyze clonal data were sensitive to clonal dispersion and to down-sampling of data. CoSpar also successfully predicted early fate biases in these datasets using only clonal information from the last time point. When clonal data was removed entirely, results were sensitive to the choice of distance metric, and no single approach optimally inferred fate bias across all data sets.

The robustness of CoSpar could greatly simplify the design of LT-scSeq experiments, by enabling experiments with fewer cells, fewer clones, or fewer time points. In all three datasets considered here, CoSpar reveals clear early fate boundaries that were not previously reported, yet in agreement with the heterogeneity of key transcription factors and fate determinants. We predicted novel transcription factors and markers in each case, and they could facilitate enriching and manipulating the desired fate outcomes.

The examples we have analyzed specifically relate to LT-scSeq implemented using LARRY^13,23^ and CellTagging^14^, but CoSpar is not limited to these technologies. The state measurement can be transcriptomic (via scRNA-seq or RNA fluorescence in situ hybridization (FISH)^38^), as shown above, as well as proteomic and epigenomic; and lineage tracing can be achieved with static DNA barcodes^13,23^, endogenous mutations^39^, or exogenous DNA constructs that accumulate mutations over time, like CRISPR-based editing^2,17,18,40,41^. CoSpar can thus facilitate interpretation of the rapidly evolving field of LT-scSeq, and thus accelerate exploration of development and disease.

CoSpar also has limitations, which directly follow from its central assumption. By enforcing coherent fate choices between similar cells (Fig. 2**a**), CoSpar becomes sensitive to choices in measuring cell-cell similarity, and to the degree of smoothing used in implementing the algorithm (Supplementary Fig. 2**c**). Thus, CoSpar will fail to identify fate biases when heterogeneity relevant to cell fate is not measured, or when it is filtered out during data analysis, or due to over- or under-smoothing. In addition, when inferring progenitor bias from clones observed at a single late time point, CoSpar necessarily leans more strongly on state information, and it might fail when heterogeneity in the later population cannot be related to heterogeneity in the initial population. Despite these caveats, CoSpar provided sensible predictions in the cases examined here.

Coherent sparse optimization could prove useful for applications beyond dynamic inference. Several problems require learning locally coherent maps from few and noisy measurements. Such problems occur, for example, when integrating two sets of measurements in the same system^42,43^ (batch correction and multi-omics), decoding spatial transcriptomes from composite FISH measurements^44^, and inferring responses of a system to individual perturbations from composite perturbation readouts^45–47^. Outside of biology, the association of measurements in one modality with sparse measurements in another can occur in marketing and social networks^48^. Forcing coherence and sparsity constraints could greatly improve map inference in general, reducing the cost of data acquisition and enabling new discoveries.

## Methods

### Definitions: states, transition maps, and clones

To formalize the problem of learning biological dynamics, we first define basic terminology. The observed **state** Of a cell can include information on its transcriptome, epigenome, proteome, metabolic state, phospho-proteome, structural organization, or a combination of all of these. It may also include information on the environment of the cell, such as the transcriptome of neighboring cells, extracellular matrix composition, etc. These are quantified by a set of *n* features, *X* ∈ ℝ^*n*^. Although *X* is continuous, it will be mathematically convenient to treat the accessible set of states as discrete. This is reasonable because experiments only sample a finite number of cells, so resolution into *X* is limited in practice. For convenience, we enumerate cell state as *X*_*i*_, or more concisely as state *i*.

In a dynamical cellular system, cells are observed to occupy a distribution of states at consecutive times, with *P*_*i*_(*t*) giving the fraction of cells in state *i* at time *t*. We consider the **finite-time transition map** *T*_*i′i*_(*t*_1_, *t*_2_) as relating between experimental timepoints through the relationship^1^:

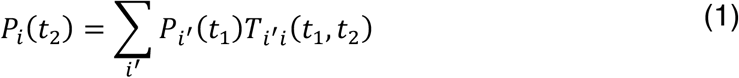

The goal of our analysis is to learn *T*_*i′i*_(*t*_1_, *t*_2_), which in turn encodes information on the fate potential of cells in each state *i*, and the rate by which cells transition between states. In typical population-sampling experiments such as scRNA-seq, the transition map is shaped by the dynamics of cells, and by the rates of cell division and loss from the tissue (see Supplementary Note 1; Supplementary Fig. 1**d**). Errors in lineage tracing affect how well we can recover the transition map (see Supplementary Note 2).

Previous work has sought to infer *T*_*i′i*_(*t*_1_, *t*_2_), from *P*_*i′*_(*t*_1_), *P*_*i*_(*t*_2_) only^1^. Here we greatly constrain the inference problem using the dynamics of clones. By **clone** We mean a set of cell states (≥ 0 cells) that arise from a common ancestor cell. Experimentally, we use “clone” to mean a set of (≥ 1) cell states that share the same barcode, a genetically heritable element. Clones may be labeled by a static barcode, or by accruing barcodes through mutation or further integration events that label sub-clones. Barcode accrual allows a cell to associate with multiple detected clonal barcodes.

### Data structures

Denoting the number of cells at time *t* as *N*_*t*_, and the number of clones as *M*, we define:

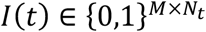: clone-by-cell matrix for the observed clonal data at time *t*, with discrete entries 0 or 1 indicating whether a cell belongs to a clone or not. We use *I*_*mi*_(*t*) to indicate its value for *m*-th clone at state *i*. For convenience, we sometimes use *i*_*t*_ to represent the matrix.
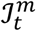: the set of cell states at time *t* that belong to *m*-th clone.
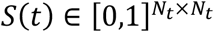: state-similarity matrix among cell states at time *t*.
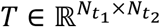: matrix of transition probability from cell states at *t*_1_ to states at *t*_2_.
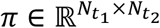: transition matrix that only allows intra-clone transitions (inter-clone transition amplitudes are set to 0).
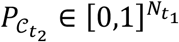: fate map, i.e., a vector of probability for each initial cell state to transition to cluster 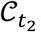 at time *t*_2_.

### Dynamic inference with CoSpar

CoSpar seeks to minimize an objective function with a close connection to compressed sensing, as discussed in the main text. A heuristic, efficient algorithm implements the optimization through an iterative procedure (see main text for the objective function, and Supplementary Note 3 for its mathematical connection with compressed sensing). Referring to Fig. 2**a**, in each iteration, we 1) threshold the map to promote sparsity; 2) enforce clonal constraints by setting inter-clone transitions to be zero and performing clone-wise normalization; 3) locally average the transition map to promote coherence. These steps are described by the following pseudo-code. Full implementation and user guide are available at https://cospar.readthedocs.io.

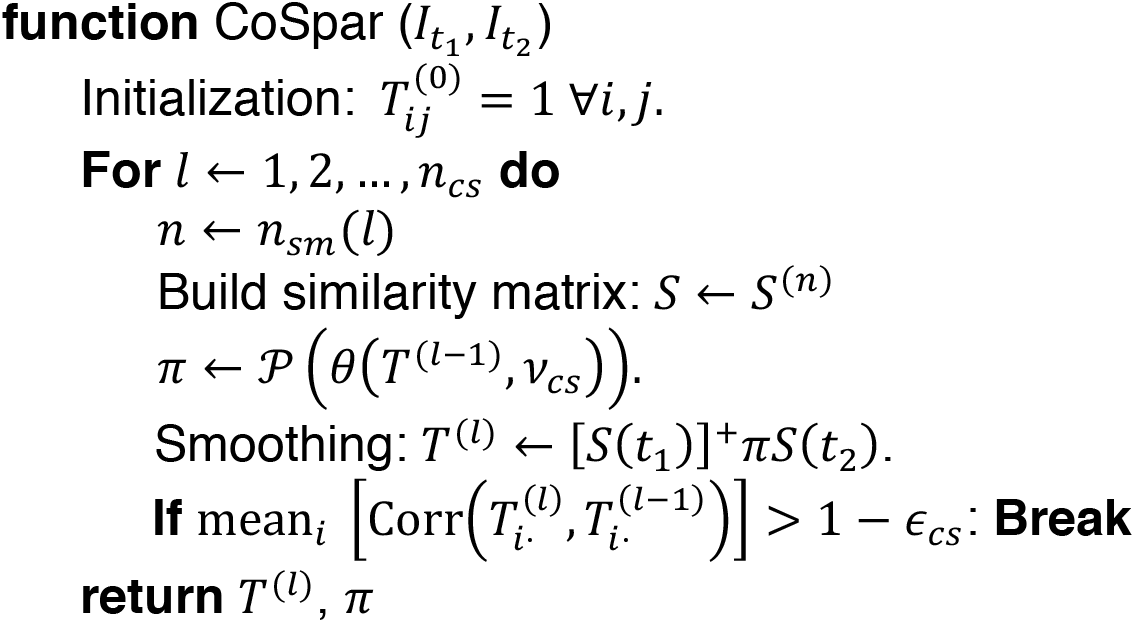

Here, + is a symbol for matrix transposition. Operators *θ*, 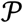 and *S*^(*n*)^ are defined below:

#### Definition of operators *θ*, 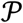

Operator *θ* implements row-wise thresholding to promote sparsity:

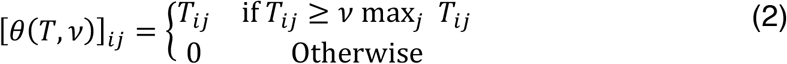

where *v* ∈ [0,1] is a parameter that tunes sparsity.

Operator 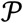 carries out clonal projection and normalization:

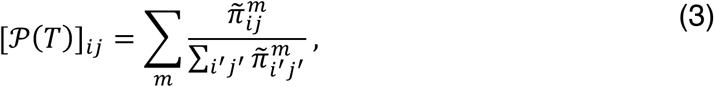

where 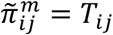 if the transition *i* → *j* occurs within clone *m*, and otherwise 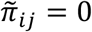. The normalization penalizes large clones, which tend to be more heterogeneous and less informative.

CoSpar has two outputs: the smoothed transition map *T* and the map *π* that only allows intra-clone transitions.

#### Similarity matrices *S*^(*n*)^

We currently know of no natural choice for establishing the similarity of two states *X*_*i*_, *X*_*j*_. We found that a Graph diffusion process^2,3^ recovered ground-truth results well in the simulations and experimental down-sampling analyses. CoSpar constructs a weighted kNN graph of observed cell states from a PCA embedding using the method proposed by UMAP^4^, leading to a graph connectivity *w*_*ij*_ from state *i* to *j* that properly takes care of the heterogeneity of local cell density, with *w*_*ii*_ = 0. To make sure that transitions between two states are reversible, we symmetrize the connectivity: 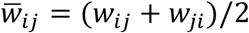. Then, the random walk matrix is

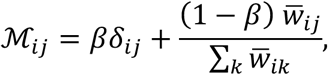

where *β* controls the probability to stay at the original state after a unit step. We then introduce a family of similarity matrices:

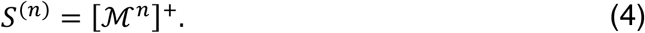

The default method implemented in *scanpy.pp.neighbors* was used to construct the kNN graph at a specified neighbor number *k*_*cs*_, with *α* = 0.1 and *k*_*cs*_ = 20.

#### Annealing steps [*n*_1_, *n*_2_, …]

CoSpar iterates through different depths *n* of *S*^(*n*)^, inspired by simulated annealing for finding the optimal solution in a rugged energy landscape^5^. Specifically, we use the sequence 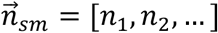 to indicate the depths at each iteration.

#### Parameter choices

The following parameters of CoSpar are adjustable: 1) parameters used for building the random walk matrices 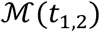, including *β* and *k*_*cs*_; 2) the sequence 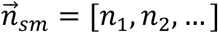 for generating annealing similarity matrix *S*^(*n*)^; 3) the threshold *v*_*cs*_ for promoting sparsity; and 4) parameters *n*_*cs*_ and *ϵ*_*cs*_ used to control iteration and convergence. We found 3 iterations are sufficient to obtain a convergent map (Supplementary Fig. 2**b,d**). Throughout this paper, we used a fixed iteration run *n*_*cs*_ = 3, and ignored *ϵ*_*cs*_ for computational efficiency. We also set *k*_*cs*_ = 20 and *β* = 0.1. We found CoSpar is more robust to *v*_*cs*_ than to 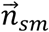 (Supplementary Fig. 2**a,c**). Other parameters are given for each respective dataset below.

### Extending CoSpar to single-time clones

When clonal data are available only at a single time point, dynamic inference is implemented as shown schematically in Fig. **2b**. Here only measurements on *I*(*t*_2_) are available. We jointly optimize the initial clonal data *I*(*t*_1_) and the transition map *T*. An iterative algorithm is used as defined here:

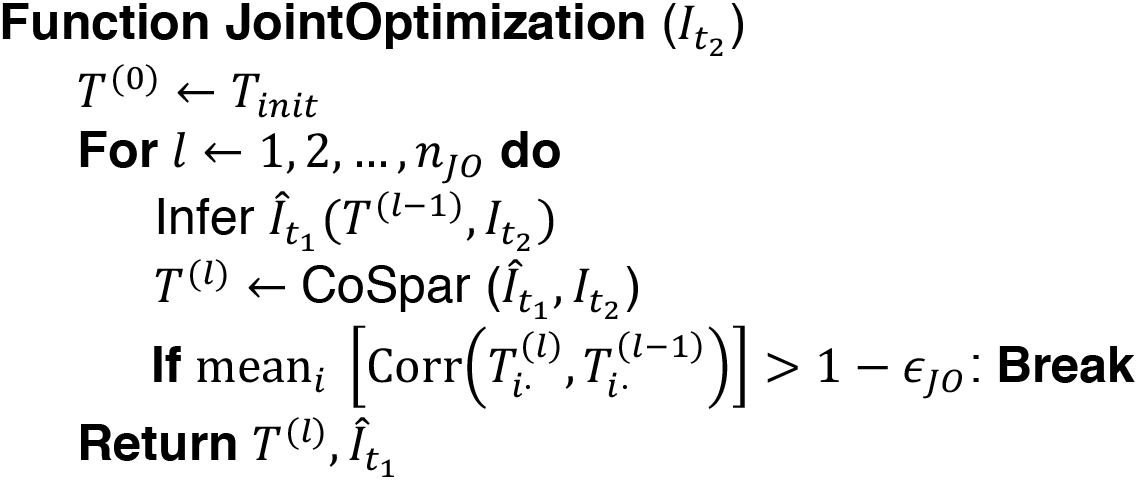

#### Initialize the map, *T*_*init*_

CoSpar uses optimal transport (OT) to construct the initialized map *T*(*t*_1_, *t*_2_) = *T*_*init*_. Given an initial state distribution at *t*_1_ and a later density at *t*_2_, OT finds a map *T*_*int*_ that minimizes the transport cost to move the initial distribution to the later one. The approach is related to that developed in Waddington-OT (WOT)^1^, but with a minor modification. To construct the OT cost matrix^1^, approximated by a cell-cell distance matrix, CoSpar offers two approaches: 1) Euclidean distance in the selected PCA space, as implemented in WOT; 2) shortest path distance on a kNN graph of the state manifold. We found that shortest-path distance generally performs better than Euclidean distance (Supplementary Fig. 3**e**; Supplementary Fig. 4**c,d**). CoSpar accepts two parameters for this initialization: a *k*_*OT*_ for constructing the kNN graph, and a regularization parameter *ϵ*_*OT*_.

#### Alternative initialization *T*_init_

OT provides a reasonable initialization when the cell-cell distance matrix contains sufficient information to match the state heterogeneity at selected time points. When this assumption fails (e.g. owing to large differentiation effects over the observed time window, or batch effects), we initialize *T* using an alternative approach, in which we generate an artificial clonal matrix based on highly variable genes at both time points: 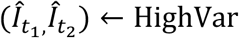, and then use it to calculate the initial transition map, 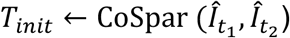. See Supplementary Note 4 for further details.

#### Inferring the clonal matrix 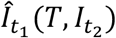

Given a transition map *T*, CoSpar updates the clonal matrix 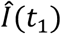 based on the principle of maximum likelihood:

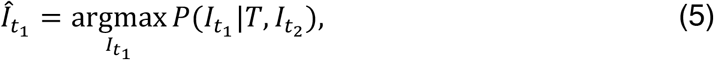

under two constraints:

1. all initial states are clonally labeled, i.e. 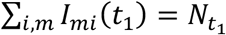;
2. the fraction of cells with a given clonal barcode structure is constant over time. Note that this constraint represents a simplification as all clones initially derive from single cells and only develop to be heterogeneous in size over time. We provide an alternative enforcing each clone to have the same size at *t*_1_, which is true for static barcoding at *t*_1_. We found that the former constraint gives robust results over all tested datasets.

These two constraints are integrated as follows. With 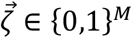 indicating a clonal barcode combination, and 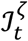 indicating the set of cell states at time *t* with barcode combination 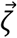, the total number of cells with the barcode structure 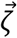 at time *t* is 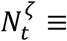 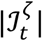. We enforce the constraint:

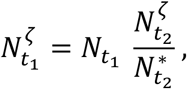

where 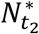 is the number of clonally labeled cells at *t*_2_. As 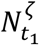 is generally non-integer, we sample the cell number probabilistically from 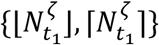, with a mean of 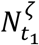, where ⌊⋅⌋ and ⌈⋅⌉ take the floor and ceil of a number, respectively.

We provide a heuristic implementation for this optimization. First, rank all observed barcode structures 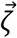 from small to large values of 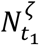. Then, sequentially infer the initial structure of each clone 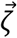:

1. compute from *T* the fate probability 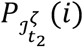 that each state *i* in *t*_1_ transitions to 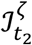, as defined below by Eq. (6);
2. select among not-yet-clonally-labeled cell states at *t*_1_ the top 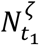 most likely initial cell states as the hypothetical initial states for this clone, and update the clonal matrix 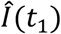 accordingly.

#### Parameter choices

The joint optimization accepts additional parameters 1) for initializing *T* (*k*_*OT*_ and *ϵ*_*OT*_ for the OT method, and gene selection parameter HighVar_gene_pctl for the HighVar method); and 2) for controlling iteration and convergence, i.e., *n*_*JO*_ and *ϵ*_*cs*_. We found that one iteration is sufficient to obtain a convergent map for all tested datasets in this paper (Supplementary Fig. 2**e**). We set *k*_*OT*_ = 5, *ϵ*_*OT*_ = 0.02, *n*_*JO*_ = 1 and ignored *ϵ*_*JO*_ throughout this paper. The remaining parameters are provided for each dataset below.

### Toolkits for transition map analysis

#### Fate map

From a transition map *T*, we can compute the probability for early states to enter a given set of states 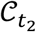 (a fate cluster). This is a key output of CoSpar, and will be used to generate other important outputs including progenitor probabilities, fate boundary, and fate coupling, etc. We first row-normalize the transition map: 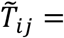 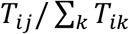. The fate probability for an initial cell state *i* is given by

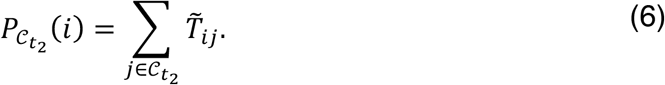

The fate probability satisfies 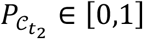.

#### Progenitor map

We compute the probability that a set of later states 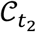 originate from a given initial state by normalizing the fate probabilities 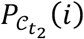 towards the fate cluster 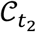:

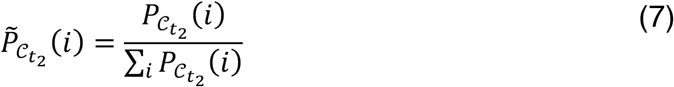

The progenitor probability satisfies 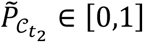.

#### Progenitor bias

We compute the bias by which an early state contributes differently to two fate clusters. Given two progenitor maps 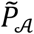 and 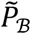 towards cluster 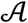 and 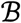, we compute the bias as

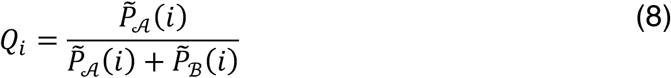

The progenitor bias is within the range [0,1]. We set state *i* to have a neutral bias *Q*_*i*_=0.5, if it a small contribution to both fates: 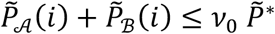, where 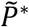 is the maximum progenitor probability across both fates, i.e., 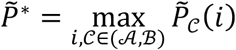. We set *v*_0_ = 0.05 in this paper.

#### Predictive genes

We perform differential gene expression (DGE) analysis between cells with different progenitor biases. The biased population towards fate 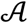 or 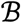 are given by

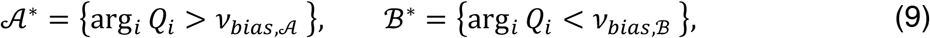

where 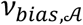 and 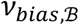 are the corresponding thresholds. We perform DGE analysis between these two populations using the Wilcoxon rank-sum test with Benjamini-Hochberg correction. We rank the enriched genes (FDR<0.05) according to the expression fold change between population 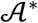 and 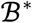.

#### Fate coupling (Supplementary Fig. 3d,f)

We define fate coupling as the correlation of fate maps towards two fates. Specifically, we first compute the fate map 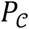 towards selected fate clusters. 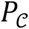 is a 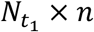 matrix where *n* is the number of selected fates, represented by cell sets 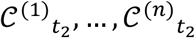. The raw coupling is given by

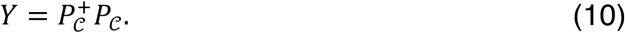

Here, *Y*_ll′_ sums over “joint probability” between fate cluster *l* and *l*′ across all initial states. We normalize the coupling as 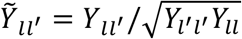, which brings the self-coupling 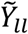 to 1, and 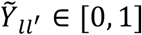.

#### Clonal fate bias (Fig. 5d; Fig. 6b)

We evaluate the fate bias of a clone towards/against a given cluster as in^6^ by quantifying the statistical significance of a clone’s occupancy of a given transcriptional state (e.g. a cluster), when compared to that expected from a random sampling of cells. The P-value (or *P*_*value*_) is computed with Fisher Exact test, accounting for the clone size. We then transform it into clonal fate bias − log_10_ *P*_*value*_, and rank each clone accordingly. We also provide the same rank plot for randomly sampled clones.

### Analyzing simulated datasets

#### Linear differentiation (Fig. 3a-d, Supplementary Fig. 2a-c)

A cell trajectory was parameterized as a one-dimensional interval of length *L*. The dynamics were simulated with a homogenous transition map corresponding to a biased random walk 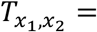 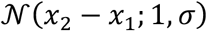, where 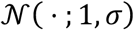 is the Gaussian distribution with mean 1 and standard deviation *σ*. Specifically, clones were simulated from this map by sampling *x*_1_~Uniform(0, L), and then *x*_2_ = *x*_1_ + 1 + *ξ* with *ξ*~ Gaussian(0,*σ*). Each pair (*x*_1_, *x*_2_) defines a clone. A total of *N* clones were simulated. To simulate barcode homoplasy, clones were randomly mixed to give *M<N* clonal barcodes of uniform size. All observations of cell states were embedded in a 50-dimensional space *Z* = (*z*_1_, …, *z*_50_) by setting *z*_1_ = *x*, and adding independent Gaussian noise *z*_*k*_ = 0.2*ξ* to each of the remaining 49 dimensions. We used *σ* = 0.5, *L* = 100, *N* = 1000. The number of detected clonal barcodes *M* was variable as shown in the figure panels. CoSpar was applied with *v*_*cs*_ = 0.2, 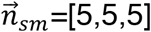.

#### Bifurcation and cell sampling (Fig. 3e-i)

A cell trajectory was parameterized as a one-dimensional interval of length *L*/2 bifurcating into two one-dimensional intervals of further length *L*/2 corresponding to fates A and B. To simulate a clonal resampling experiment, for each clone an initial barcoded cell was seeded at *x*_0_~Uniform(0, L) at *t* = 0. Cells were simulated to divide once at each unit time step, and all cells progressed along the trajectory according to a random walk, with 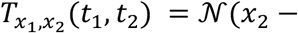 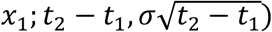. As each cell transitions past the bifurcation point (L/2) it chose between fates A, B with probability 1/2. At *t* = *t*_1_, we sampled cell states in each clone with a success rate 0.5 per cell. Successfully sampled cells were removed, and the remaining unobserved cells continued to divide and progress as described. The state of all remaining cells was profiled at *t*_2_ = *t*_1_ + 1. The observed cell states were embedded in a 50-dimensional observation space *Z* by first embedding in two-dimensions,

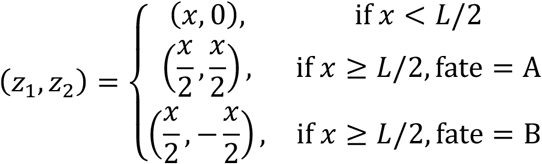

and then adding independent Gaussian noise *z*_*k*_ = 0.2*ξ* to each of the remaining 48 dimensions. We set *σ* = 1, *t*_1_ = 5, *L* = 10. *M*=100 clones were simulated. CoSpar was applied with *v*_*cs*_ = 0.2, 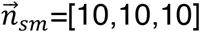.

#### Evaluating CoSpar with simulated data

We defined the TPR (Fig. 3**d,g**) as the fraction of rows of the inferred transition map, 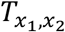 for which the maximum transition rate is within 3*σ* of the expected peak position, i.e. 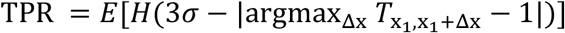 where *E*(·) is the mean over all rows of T, and *H*(*z*)={1 for *z*>0; 0 otherwise}. The progenitor bias for the bifurcation model _(Fig. 3**h,i**)_ was calculated according to Eq. (8). Each of the TPR and progenitor bias comparisons (Fig. 3**d,g,i**) shows averages after application of CoSpar to 5 independent simulations.

### Benchmarking and applying CoSpar to hematopoiesis

#### Pre-processing

Data^7^ is available at Gene Expression Omnibus (GEO), accession number GSE140802. Data was preprocessed as originally described^7^: 1) UMI counts were normalized in each cell to the average across all cells; 2) highly variable genes were selected using the SPRING gene filtering function (filter_genes using parameters *min_vscore_pctl* =85, *min_counts*=3, *min_cells*=3)^8^; and 3) genes correlated with cell cycle were excluded from the highly-variable gene list (genes with correlation *C* > 0.1 to the signature genes defined by *Ube2c, Hmgb2, Hmgn2, Tuba1b, Ccnb1, Tubb5, Top2a*, and *Tubb4b*). The 2-dimensional embedding and state annotation of cells were as in^7^, also available at the GEO website (GSE140802). We selected the top 40 Principal Components (PCs). Unless otherwise stated, we constructed kNN graph with *k* = 20 for downstream analysis.

#### Applying CoSpar

Code detailing implementation of CoSpar to the data is provided at https://cospar.readthedocs.io/. In brief, we evaluated the progenitor fate bias, identified putative driver genes, and computed the fate coupling as described above. The default parameters are *v*_*cs*_ = 0.1, 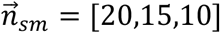, and we initialize the transition map using the OT method for joint optimization.

#### Intra-clone dispersion (Fig. 4b)

We quantified the intra-clone dispersion of a clone *m* as the maximum cell-cell distance *d*(*m*, *t*) within a clone at time *t* (*t* = 2, 4, 6), where the distance was measured by the shortest-path distance in the kNN graph at *k* = 5. Fig. 4**b** Shows the dispersion normalized by the mean dispersion on day 2.

#### Transition map using the method from Weinreb et al^[Citation error]^ (Fig. 4c,h; Supplementary Fig. 3a,b, g-i)

We selected clones that have a unique fate at a later time point, where each mature fate cluster was defined as in Weinreb et al (see annotations at Fig. 4**a**). Multi-fate clones were discarded. Given this clone matrix *I*^w^(*t*), with *t* = 2,4,6, we computed the transition map as 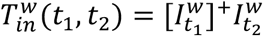, where any initial cell state has the same probability to transition to any later cell state observed in the same clone. The ground truth progenitor bias in Fig. 4**c** Shows the progenitor bias *Q*_i_ on day 2 and day 4 computed from 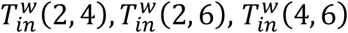 using Eq. (8).

#### Fate map reconstruction error (Supplementary Fig. 3a,b)

To allow comparison between methods, we used *π*(4, 6) from CoSpar with *v*_*cs*_ = 0.2 or 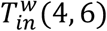 from the Weinreb method, constructed from sub-sampled clones on day 4-6, to compute the fate map 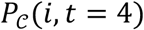 towards cells annotated with a given fate (cell set 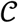) according to Eq. (6). We evaluated the inferred maps by comparing them to a ground-truth fate map 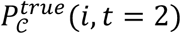 from the Weinreb method with all clones from day 2-4. We evaluated the prediction using the Wasserstein distance^9^ between the two distribution 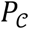 and 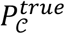, restricted to the progenitor state space 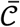 (i.e., excluding states belonging to fate 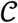). Note that 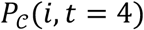 maps the fate probability of cells sampled on day 4, while 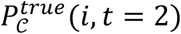 is for cells sampled on day 2. To compare the fate maps for these non-overlapping cell subsets, we computed the OT map *T*^*OT*^ from day-2 states to day-4 states with *k*_*OT*_ = 5 and *ϵ*_*OT*_ = 0.02, using shortest-path distance. The Wasserstein distance is given by 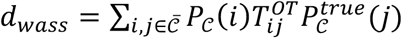. We computed the Wasserstein distance for 3 major fates: Neutrophils, Monocytes, and Basophils, and reported the average.

#### Waddington-OT (Supplementary Fig. 3f; Supplementary Fig. 4e)

Results shown were obtained using the WOT package (https://github.com/broadinstitute/wot)^1^, using default parameters: *ϵ*_*OT*_ = 0.05, *λ*_1_ = 1, *λ*_2_ = 50.

### Benchmarking and applying CoSpar to fibroblast reprogramming

#### Pre-processing

Data was downloaded from GEO, accession number GSE99915. We followed the same processing as described above for hematopoiesis, and removed cell-cycle-correlated genes with correlation score |*C*| > 0.03. We used UMAP (scanpy.tl.umap with *min_dist*=0.3) to generate the embedding.

In this dataset, cells were barcoded at three time points (day 0, 3, and 13). Following Biddy et.al.^6^, we concatenated day-0 and day-3 barcodes to form a unique clonal ID for downstream analysis. However, keeping 3 barcodes per cell, thus allowing nested clonal structure, works equally well (Supplementary Fig. 4**f-h**). We also inherited their annotation for the reprogrammed cluster (obtained by email communication with the authors), and used their selected clones to define the ground truth for reprogramming and failed trajectories. The failed cluster (Fig. 5**a**) was defined as a leiden cluster (scanpy.tl.leiden with *resolution*=1.5) in the cells sampled at day 28, which highly expresses *Col1a2* (Supplementary Fig. 4**a**), a gene expressed in fibroblasts that failed reprogramming^6^. The reprogrammed and failed cluster were used to define the progenitor bias in this dataset.

#### Applying CoSpar

The default parameters are *v*_*cs*_ = 0.2, 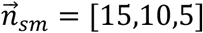, and we initialize the transition map using the OT method for joint optimization. See jupyter notebook implementation at https://cospar.readthedocs.io/.

#### Selecting dispersed clones (Fig. 5d,e)

We first calculated for each clone the fraction *γ*_*o*_ of cells within the reprogrammed cluster. Dispersed clones are defined as occupying both the reprogrammed cluster and other states on day 28, thus having intermediate values of *γ*_*o*_. We selected dispersed clones satisfying *R*_−_ ≤ *γ*_*o*_ < *R*_+_, where *R*_−_ = *x* and *R*_+_ = 0.4 − 2*x*, and *x* parameterizes the window. This parameterization was chosen so that we could evenly exclude clones at both sides of the window when adjusting *x*. The fraction of clones within this window was used as an indicator for each sub-sampled dataset in Fig. 5**e**.

#### Transitions using the method from Biddy et al (Fig. 5e,f)

Following Biddy et.al.^6^, we first identified clones that are enriched or depleted in the reprogrammed cluster according to Fisher’s Exact test. Among statistically significant clones (*P*_*value*_ ≤ 0.05), we selected cell states belonging to reprogramming clones (*γ*_*o*_ > 0.4) as putative reprogramming population 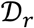, and classified cell states of low-reprogramming clones (*γ*_*o*_ < 0.4) as putative failed population 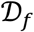.

To boost the performance for downstream analysis, we made the following modification to the original method in Biddy et.al.^6^. For a putative population (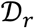 or 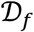), we enriched for high-fidelity states by iteratively excluding clones with *γ*_*o*_ closest to 0.4 until the total number of cells in 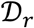 or 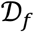 was at or below 3,000.

#### Calculating marker gene TPR (Fig. 5e,f, Supplementary Fig. 4b)

For a putative reprogramming 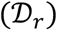 and failed 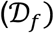 population predicted by either CoSpar or the Biddy method, we assessed their accuracy by the overlap of their top differentially expressed genes with those from the reference population (defined by the fate-biased clones selected by Biddy et.al.^6^).

To predict population 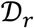 and 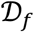 with CoSpar, we inferred *T* with *v*_*cs*_ = 0.4 and threshold the fate map 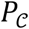 built from the intra-clone transition map 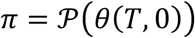 as follows:

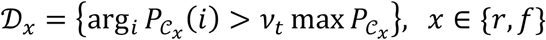

where, to enrich for high-fidelity states, *v*_*t*_ = max (0.5, *ω*) and *ω* was chosen such that 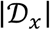 is the largest value below 500.

For both CoSpar and the Biddy prediction, when 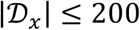, we increased the total cell number up to 200 by adding the nearest neighbors of selected cell states using the kNN graph defined by the full dataset. This step supports the statistical power of the differential gene expression (DGE) analysis.

Finally, we performed DGE analysis between 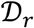 and 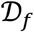, identified enriched genes for each population, and compared them with the reference. Specifically, we first calculated the P-value for each gene using the Wilcoxon rank-sum test, with Benjamini-Hochberg correction. We ranked them according to the expression fold change between 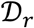 and 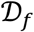, kept the top 50 genes enriched in 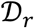 and another top 50 in 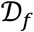, and excluded statistically insignificant ones (adjusted P-value ≥ 0.05). Denoting the resulting gene set for predicted population 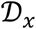 as *ε*_*x*_, and that from the corresponding reference population as 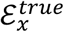, the marker gene TPR for this putative population is given by

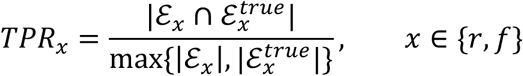

The final marker gene TPR for a given method (CoSpar or the Biddy method) was (*TPR*_r_ + *TPR*_*f*_)/2.

### Application of CoSpar to in vitro differentiation of lung endoderm

#### Pre-processing

Data was downloaded from GEO, accession numbers GSE137805 and GSE137811. We selected highly variable genes using filter_genes function (*min_vscore_pctl*=80, *min_counts*=3, *min_cells*=3), and normalized the UMI counts per cell to 10000. We used the top 40 PCs to construct kNN graph with *k* = 20 for downstream analysis. We inherited the original embedding on day 17 and 21 by Hurley et.al.^10^ (available at https://kleintools.hms.harvard.edu/tools/springViewer_1_6_dev.html?cgi-bin/client_datasets/nacho_springplot/allMerged), and used UMAP (scanpy.tl.umap with *min_dist*=0.3) to generate the embedding for day-15 and day-27 cells. The iAEC2 cluster is defined as the day-27 leiden cluster (scanpy.tl.leiden with *resolution*=0.5) that highly express *SFTPB* and *SFTPC* (Supplementary Fig. 5**a**), marker genes for iAEC2 cells^10^.

#### Applying CoSpar

To apply joint optimization (Fig. 6**c**; Supplementary Fig. 5**f,g**), we initialized the transition map using the HighVar method with *HighVar_gene_pctl*=80, and ran CoSpar with *v*_*cs*_ = 0.2, 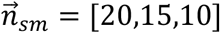. See jupyter notebook implementation at https://cospar.readthedocs.io/.

## Data availability

All data analyzed in this article are publicly available through online sources. The annotated data, results, and Python implementation are available at https://cospar.readthedocs.io/. The raw data for the hematopoiesis dataset can be accessed at Gene Expression Omnibus (GEO) database with accession number GSE140802, the reprogramming dataset via GSE99915, and the lung dataset with GSE137805 and GSE137811.

## Code availability

The results reported in this paper and our Python implementation are available at https://cospar.readthedocs.io/.

## Acknowledgements

SWW acknowledges support by Damon Runyon Computational Biology Fellowship. AMK acknowledges support by NIH Grant 1R01HL14102-01. We thank Tal Scully for helping with figures. We thank Darrell Kotton, Killian Hurley, and Michael J Herriges for feedback and support with AEC2 analysis, and for comments on the manuscript.

## Author contributions

SWW and AMK conceived the project. SWW devised the computational method, wrote the package, and carried out CoSpar analyses. SWW and AMK wrote the manuscript. AMK supervised the project.

## Supplementary Information for

**Supplementary Fig. 1.**
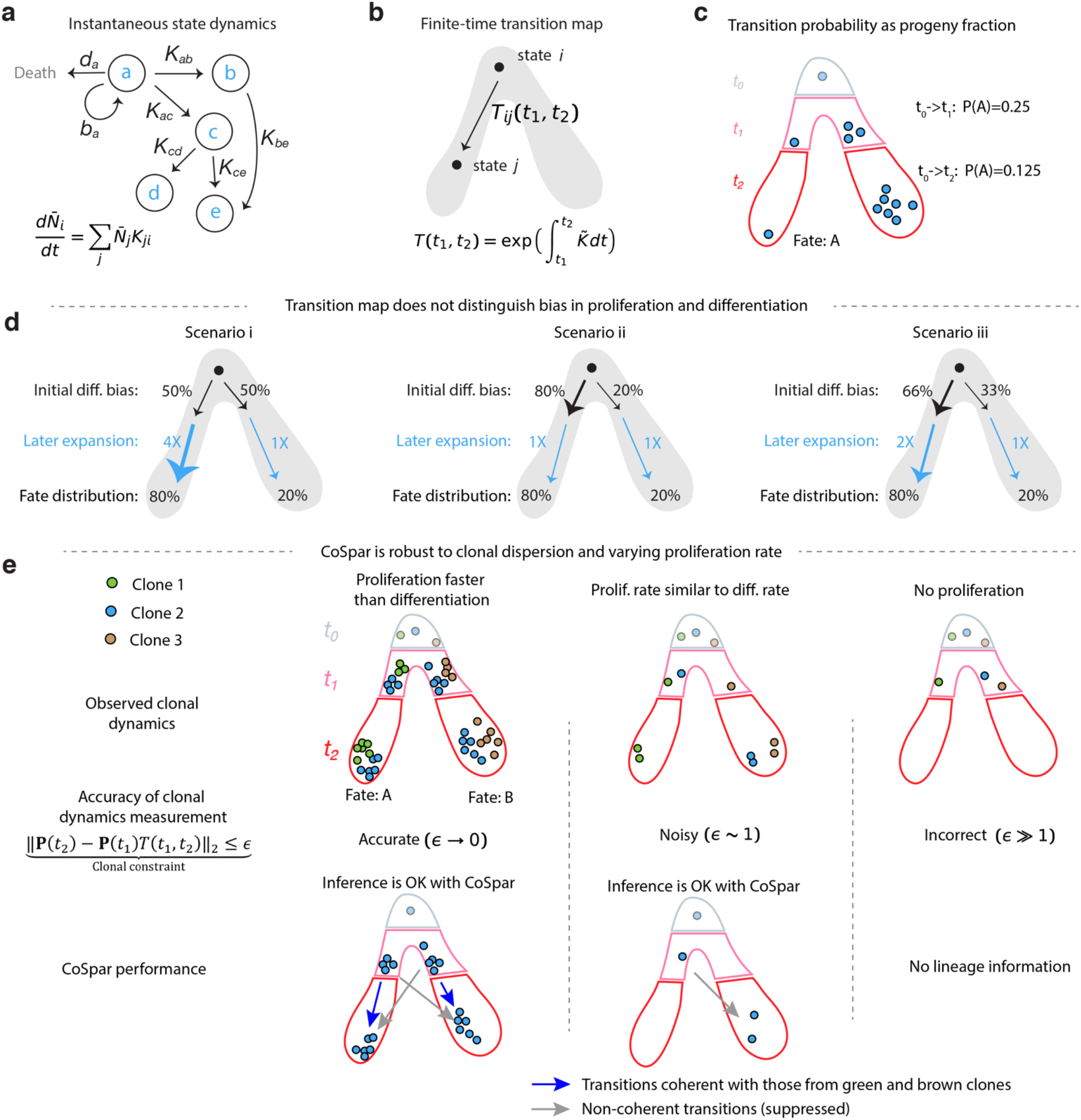
Models, assumptions and limitations of Coherent Sparse Optimization. **a**, Simple example of the class of stochastic models that CoSpar seeks to learn. In such models, each node represents an observed cell state. In practice, thousands of measured states are included; here only five are shown. At each state cells self-renew, die, or differentiate with state-specific rates. The mean fraction of cells in each state evolves according to coupled first-order equations as shown. See Supplementary Note 1 for details. **b**, The empirically-observed finite-time transition map can be interpreted through its relation to the transition rate matrix *K* (see panel **a**). See Supplementary Note 1 for details. **c**, Schematics illustrating the operational, experimentally-accessible definition of a transition probability, as the average fraction of progeny derived from an initial cell *i* at *t*0 that differentiates into a target state *j* at later times. As defined, transition probabilities are sensitive to biases in fate choice, and to differential rates of cell division and cell loss. **d**, Schematics exemplifying that transition maps cannot distinguish fate bias from differences in net rates of cell expansion (division – loss). Three different underlying dynamics lead to the same transition maps. **e**, Schematics clarifying the robustness of CoSpar to clonal dispersion (demonstrated in Fig. 3). i), When cells undergo extensive proliferation prior to fate bifurcation and clonal sampling, each clone densely samples several differentiation trajectories. By imposing sparsity and coherence, CoSpar re-enforces a minimal number of transitions that explain dynamics across all clones. ii), At lower rates of proliferation, fewer cells from each clone are sampled, and it may lead to observing clonally-related cells at different time-points on different trajectories, as shown (blue clone sampled towards fate A at *t*_1_, and towards fate B at *t*_2_). By enforcing coherence between clones rooted in neighboring states, CoSpar may still recover a correct transition map. In this case, there is a trade-off in the CoSpar cost function between minimizing the clone transition map error and maximizing coherence. iii), Lacking proliferation, one cannot establish clonal relationships that constrain dynamic inference.

**Supplementary Fig. 2.**
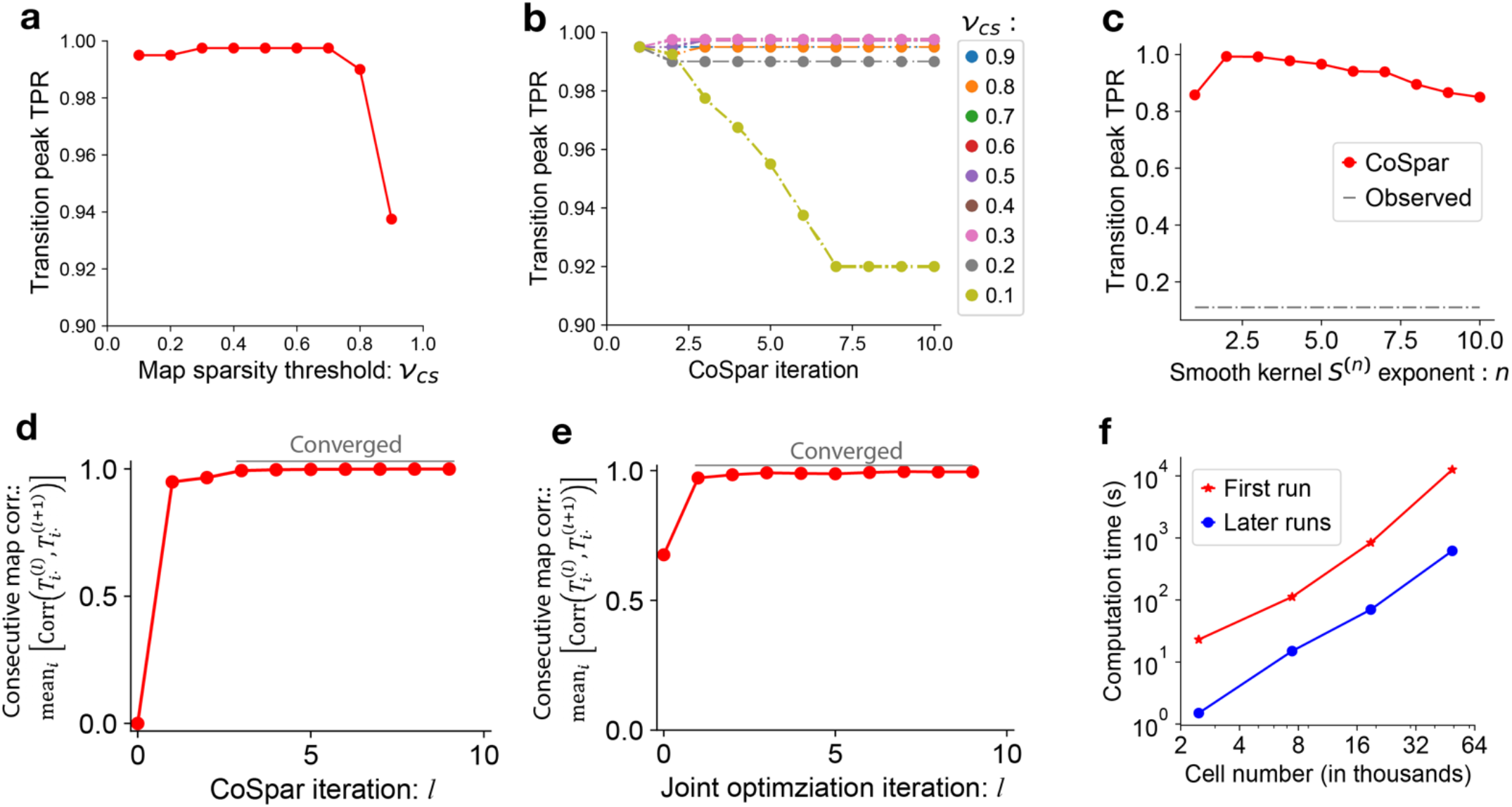
Evaluating CoSpar performance across parameter sweeps. **a-c**, Performance of CoSpar using simulations as in Fig. 3**a-d** With a range of algorithm parameters (see Methods for parameter definitions): (**a**) sparsity threshold *v*_*cs*_ ∈ [0,1]; (**b**) number of iterations, showing convergence; (**c**) smoothing kernel exponent. **d,e**, Demonstration of algorithm convergence, seen in the correlation between maps from consecutive iterations against the number of iterations, for the two algorithms (CoSpar, and Joint CoSpar, see Methods). The maps analyzed here correspond to those from the down-sampled hematopoietic dynamics (Fig. 4**h**). **f**, Computational time to convergence, as a function of total cell number. In the first run, CoSpar will generate (and save) a similarity matrix, which is very costly (red curve). CoSpar can use similarity matrices generated previously to speed up computation (blue curve).

**Supplementary Fig. 3.**
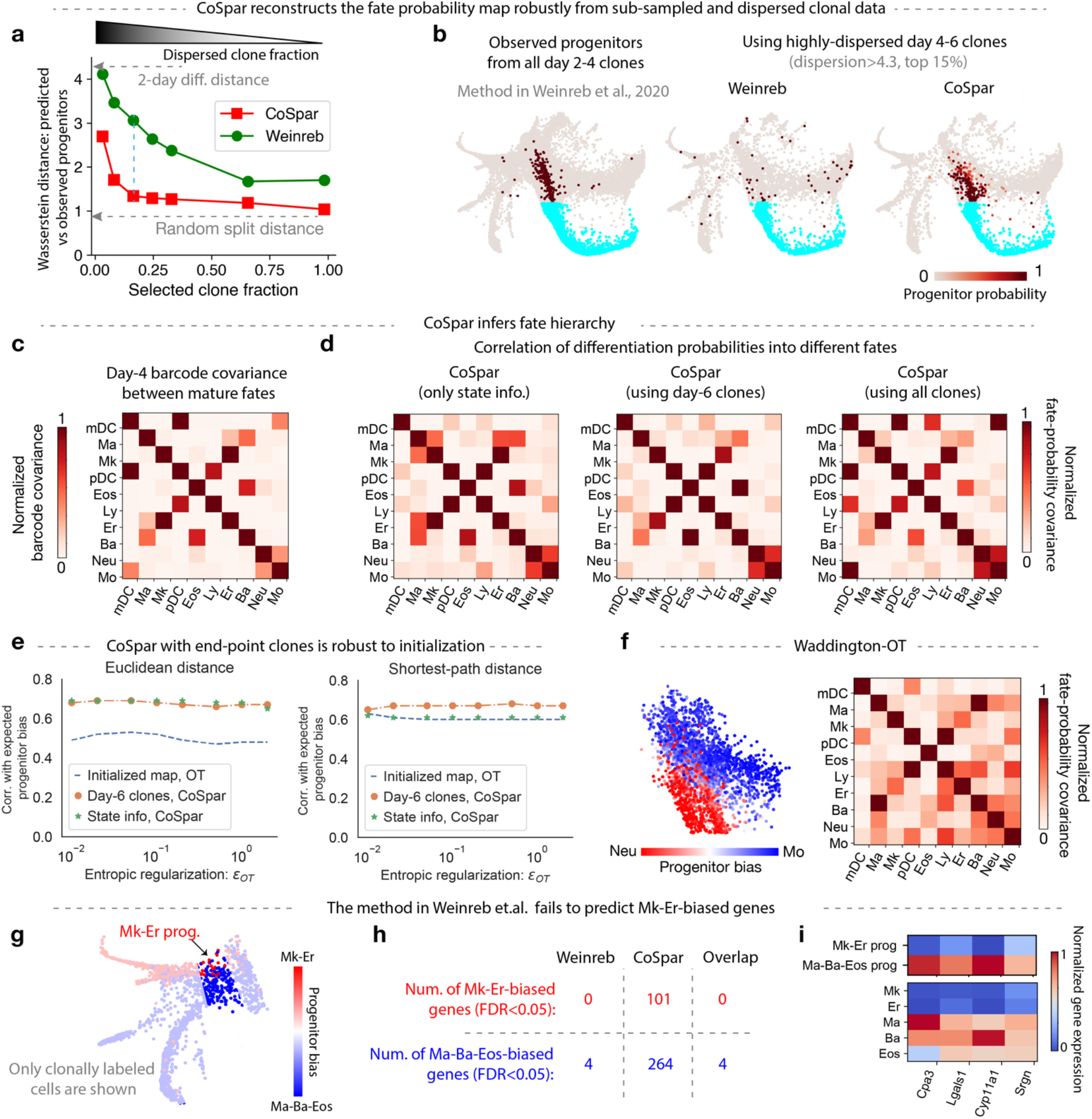
Benchmarking CoSpar in hematopoiesis. **a**, CoSpar reconstructs transition maps from sub-sampled and dispersed clonal data. Here, we evaluate the prediction error as the Wasserstein distance between fraction of cell progeny predicted to occupy a given fate, compared to that obtained from the ‘ground truth’ transition map constructed using all clonal data rooted in day 2 clones (see main text). In **a**, the prediction error is assessed for a decreasing fraction of day 4-6 clones, obtained by progressively excluding less dispersed clones that contribute the strongest signal (see Fig. 4**b**). Green curve is obtained by applying the method from the original paper. A lower bound on the error (random split distance) is the Wasserstein distance between random 50% partitions of the ground-truth data. The largest observed errors are comparable to the Wasserstein distance between populations separated by two days of progressive differentiation (upper grey arrow). **b**, The ground truth and predicted fate maps for neutrophils cluster using the 15% most dispersed clones. These plots illustrate one value on the plot in **a**. **c**, The normalized covariance of clonal barcode abundances between different cell types, calculated using all data on day 4 of differentiation^1^. **d**, The correlation of predicted transition probabilities of progenitors, inferred with CoSpar using different data indicated (See Methods). **e**, Joint CoSpar optimization is robust to initialization and choice of distance metric. This panel accompanies Fig. 4**g**. Plots show the correlation of progenitor biases calculated from the transition maps for different initialization choices of the transition map. Optimal transport (OT) is used to initialize the transition map from state information alone prior to CoSpar. Plots scan the OT entropic regularization strength *ϵ*_*OT*_. **f**, Application of Waddington-OT (WOT) to hematopoiesis dataset. WOT was applied to the same data in Ref^2^, where clonal data was used to tune the local cell proliferation rates. When WOT is applied without access to any clonal information, performance is degraded as seen by comparing the plots here to the ground truth. Plots are to be compared with those in panels **c**,**d** And Fig. 4**c**. WOT is applied with default parameters (*ϵ*_*OT*_ =0.05). **g-i**, Predicting early fate boundaries in the Gata1+ lineages using the original method from Ref^2^. **g**, Predicted progenitor bias among the Gata1+ cells on the state embedding. **h**, Comparison of the number of differentially expressed genes (FDR<0.05) identified from different methods of clonal analysis. **i**, Gene expression heat map for all differentially expressed genes identified with the Weinreb method^2^.

**Supplementary Fig. 4.**
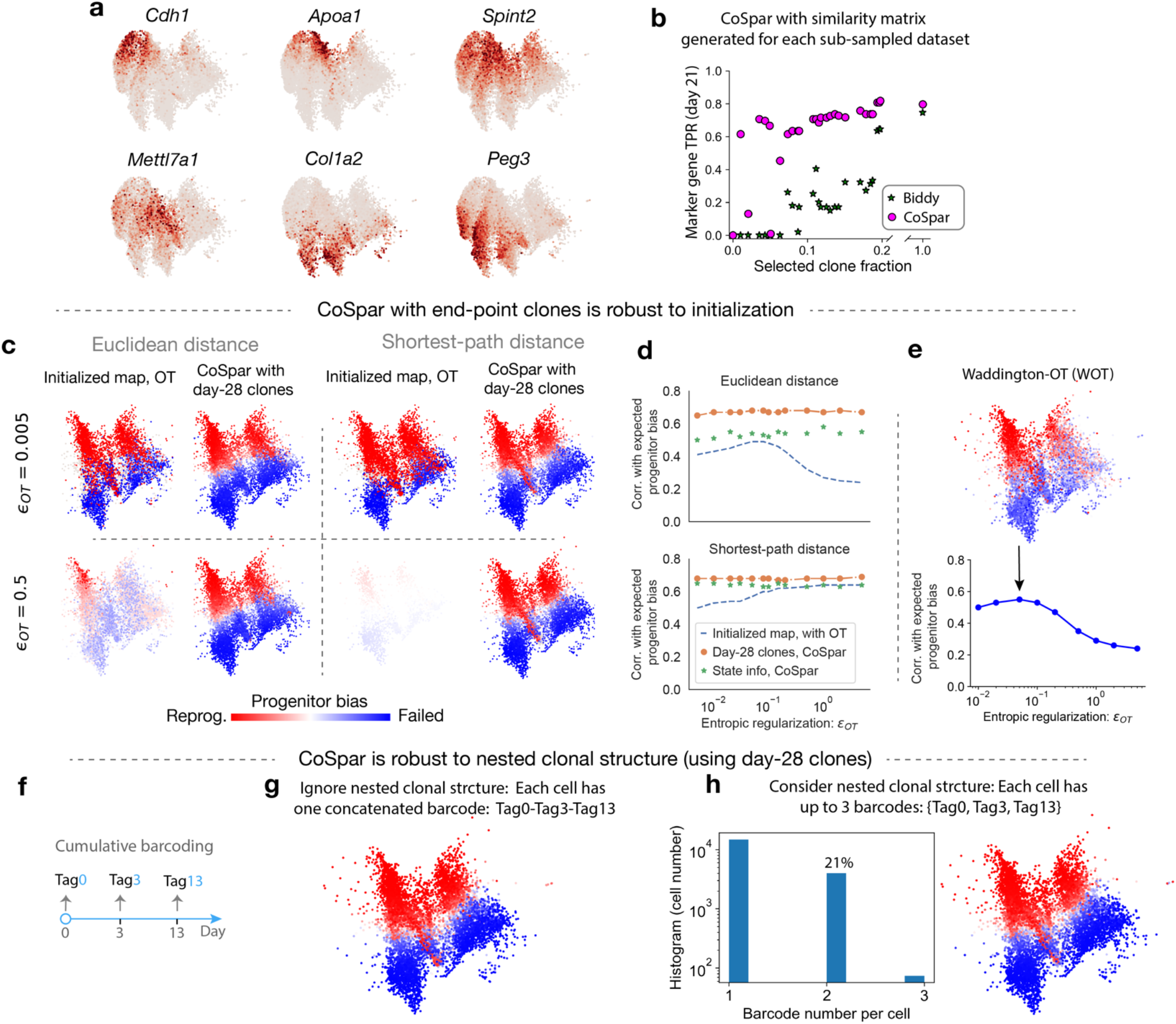
Benchmarking CoSpar in fibroblast reprogramming. **a**, Expression of selected marker genes on UMAP visualizations from day 15, 21 and 28. **b**, Reproduction of results in Fig. 5**e** Using a similarity matrix obtained from each sub-sampled dataset. Results are seen to be robust to sub-sampling strategies. **c-e**, Transition maps inferred by CoSpar with access only to end-point clonal information are robust to the choice of initialization. These panels accompany Fig. 5**h**. **c**, Visualization of the progenitor bias derived from the initialized transition map and the corresponding CoSpar prediction, for different entropic regularizations and distance metrics as indicated. **d**, Parameter sweep quantifying the stability of the predicted progenitor bias. **e**, Progenitor bias prediction from Waddington-OT^3^, which relies only on state information. Upper panel: the predicted progenitor bias on the state manifold at *ϵ*_*OT*_=0.05. Lower panel: progenitor bias correlation with ground truth across different *ϵ*_*OT*_ values. **f-h**, CoSpar analysis with clonal barcodes integrated at sequential time points. The analysis was done with clonal data on day 28. **f**, The cumulative barcoding scheme in the reprogramming experiment. Cells were barcoded on day 0, 3, and 13. **g**, A progenitor bias prediction generated by concatenating all tags from all three time points into a single clonal barcode for each cell, thus ignoring the nested clonal structure in the data. **h**, Equivalent results of CoSpar analysis with nested clonal structure, carried out by treating Tag0, Tag3 and Tag13 as independent barcodes for a cell, such that each cell may have up to three barcodes. Left panel shows the histogram of barcode number per cell.

**Supplementary Fig. 5.**
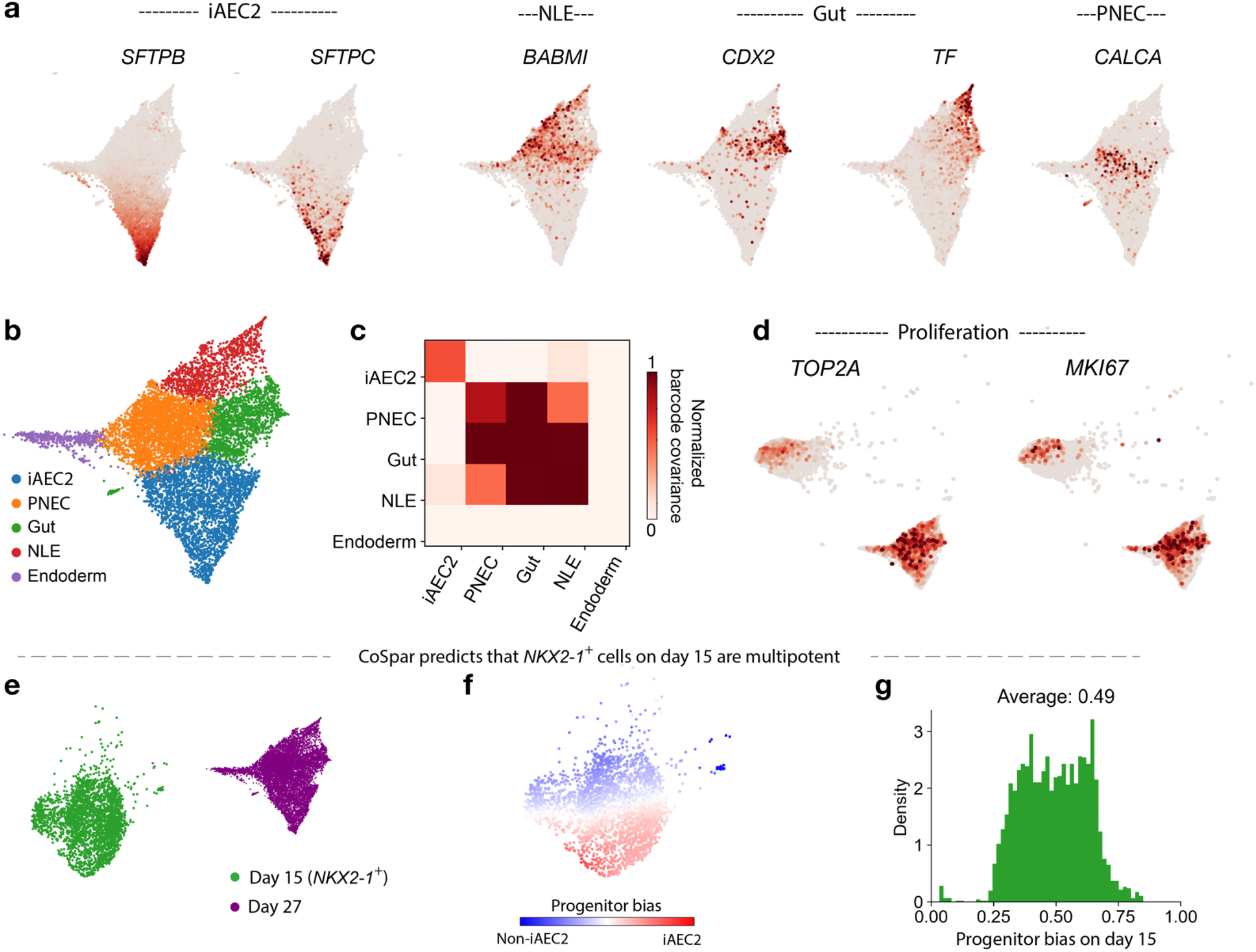
Marker gene expression and clonal structure during differentiation into alveolar cells and other endodermal cells. **a**, Expression of genes associated (in Ref^4^) with iAEC2 cells, non-lung endoderm (NLE), gut endoderm, and pulmonary neuroendocrine cells (PNEC). **b**, Leiden clustering of day-27 cell states. Cluster are named based on their corresponding gene expression. **c**, Normalized barcode covariance on day 27 among all clusters, showing evidence of clonal partitioning of iAEC2 cells. **d**, Expression of two representative genes marking proliferating cells (*TOP2A* and *MKI67*) on day 17 and 27 state manifold, showing that cells predicted by CoSpar to show low commitment on day 17 appear proliferating (Fig. 6**c**). **e-g**, CoSpar predicts that lineage restriction occurs after day 15, except for a rare fraction of cells committed to non-iAEC2 fates. **e**, UMAP visualization of cell states on day 15 and 27. **f**, CoSpar-predicted progenitor bias among cells on day 15. **g**, Histogram of the progenitor bias on day 15 (shown in panel **f**). Unlike on day 17 (Fig. 6**c**), here progenitor bias is concentrated at 50%.

**Supplementary Fig. 6.**
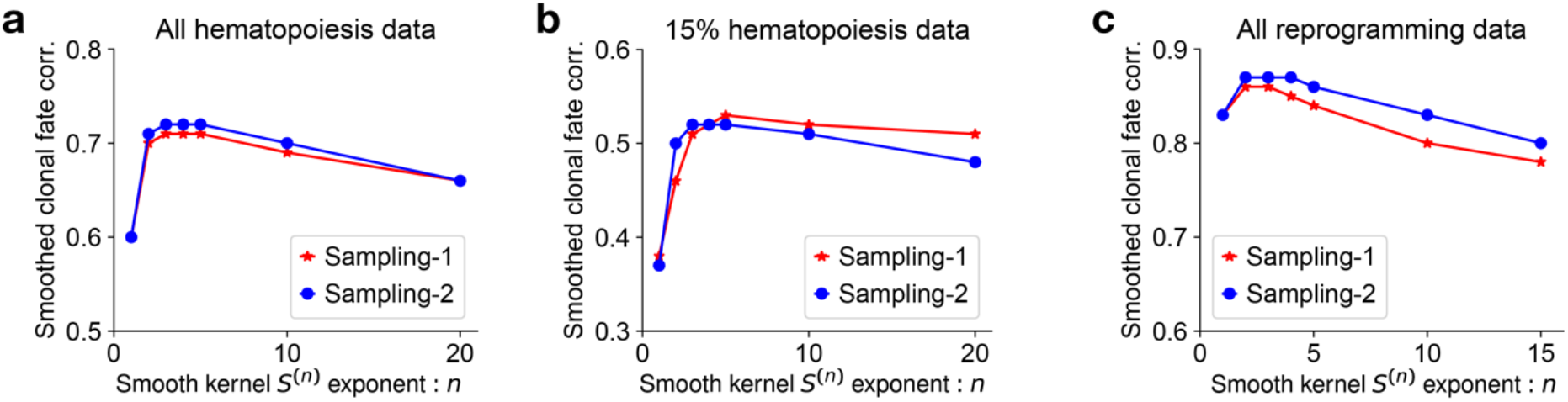
Establishing upper bounds for fate prediction after data loss. In this paper, performance of CoSpar was compared to previously published methods by discarding clonal data and then examining the fidelity of fate predictions in the face of data loss. Supporting the results reported in Figs. 4**g,i** And 5**h**, we obtain an upper bound for fate prediction, by randomly sampling 50% cells from the full ground-truth dataset in each case to predict the progenitor bias of remaining cells, with different smoothing exponents *n*. Prediction was carried out by first inferring the progenitor bias 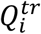 from the training data (denoted by *tr*) to predict the bias 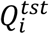 of the test data, by imputation via graph diffusion: 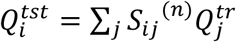. Results show that, in all the three cases considered, a smoothing exponent *n*=3 provided the best correlation between the imputed and actual values of 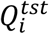. These correlation values are indicated by the upper dashed grey lines in Figs. 4**g,i** And 5**h**.

### Supplementary Note 1: Connecting transition maps to models of differentiation

This note grounds the finite-time transition map in a stochastic model of cell differentiation. In doing so it also clarifies what cannot be learnt from the transition map.

We begin by considering a Markov model of differentiation represented by an arbitrary graph of finite size, where each node represents a cell state. In this model, each cell probabilistically undergoes proliferation, death, and differentiation with rates that are specific to the cell state. A clone is a realization of such a stochastic branching process, seeded as a single barcoded cell in some cell state. Starting from a cell state *i*, *k*_*ij*_ is the differentiation rate to a different state *j*; *b*_*i*_ is the probability of a cell dividing into two cells; and *d*_*i*_ is the cell loss rate for cells in state *i*. We assume that these rates are first-order (independent of the number of cells in a state). These rates can vary with time to reflect changes in the tissue environment. Supplementary Fig. 1**a** Shows a simplified example of such a model.

This model is useful in its simplicity, but it is clearly not general: being a Markov process, it assumes that we have a complete measurement of the variables that could affect state dynamics, such as the transcriptome, epigenome, and extracellular environment. This is unlikely to be true. Incomplete state measurement leads to a non-Markovian dynamics^5^. Nonetheless, our model may be a useful approximation as it generates predictions of biomarkers and fate regulators, and their correlation with fate bias.

Our goal in this paper is to learn the structure of such a graphical model (e.g. Supplementary Fig. 1**a**) and its rate constants, from LT-scSeq data. To learn a model from data, we focus most simply on the mean dynamics of cell number at each state. To do so, one could consider a complete stochastic description using the chemical master equation^6^, which gives the distribution evolution over the extended state space *N* × *X* = {(*N*_*i*_, *X*_*i*_) ∀ *i*; and *N*_*i*_ = 1, 2, …}, where *N*_*i*_ is the number of cells at state *i* and *X*_*i*_ is the corresponding state. However, because we assume a first-order model, there exists a closed-form equation for the dynamics of average cell number 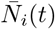 at state *i* and time *t*,

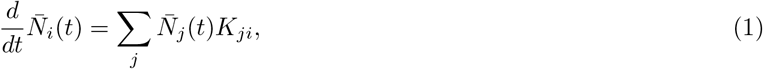

where *K*_*ij*_ ≡ (1 − *δ*_*ij*_)*k*_*ij*_ + *δ*_*ij*_(*b*_*i*_ − *d*_*i*_ − Σ_*k*≠*i*_ *k*_*ik*_), with *δ*_*ij*_ = {1 if *i* = *j*; otherwise 0}, is the instantaneous transition rate from state *i* to *j* that includes all cellular processes: division, cell death, and differentiation. This mean dynamics only captures the net effect of cell number change (*b*_*i*_ − *d*_*i*_), and does not distinguish whether it is from cell proliferation or loss.

To make contact with experiment, we represent the number of cells at each state as a fraction of the total cell number to obtain the cell density:

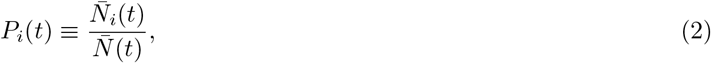

where 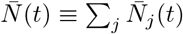 is the total cell number at time *t*. The dynamics of the cell density *P*_*i*_(*t*) is

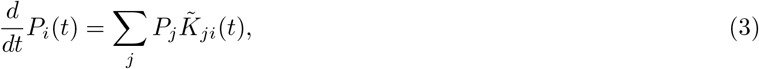

where 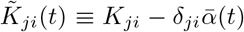, and 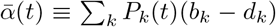 is the average growth rate of the population at time *t*. Diagonal elements in 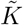 reflect whether net growth in each state is larger (positive) or smaller (negative) than the population average.

We now can ground the transition map *T* in terms of the model. Integrating Eq. (3) from time *t*_1_ to *t*_2_ leads to the relation

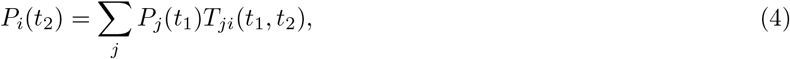

where the intrinsic finite-time transition map

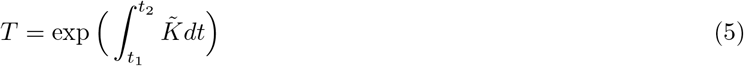

is obtained from matrix exponentiation of the corrected instantaneous transition rate matrix 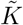.

The transition probability *T*_*ij*_ is the fraction of progenies from initial state *i* that ends at later state *j* (Supplementary Fig. 1**b**). To see this, we can sum over all states in Eq. (4), and noting that Σ_*i*_ *P*_*i*_(*t*) = 1, we have 1 = Σ_*j*_ *P*_*j*_(*t*_1_) Σ_*i*_ *T*_*ji*_. This equation is valid for any distribution *P*_*j*_(*t*_1_) and therefore the transition map satisfies the conservation property

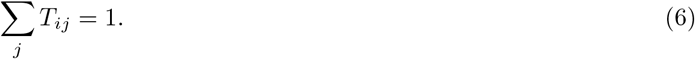

Owing to its normalization (Eq. 6), the transition map that is experimentally accessible captures the most interesting property we want: the probability of a cell to differentiate into different cell types. A certain initial state *i* can transition to multiple states over time window *t*, i.e., *T* has multiple non-zero entries associated with the *i*-th row.

Nonetheless, it is important to note that *T*_*ij*_ is shaped both by differences in transition rates between states, and by the collective effect of proliferation and cell death along the trajectories between state *i* and *j*. Mathematically, although proliferation and cell death only affect the diagonal terms in the instantaneous transition matrix 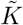, the matrix exponentiation in Eq. (5) will propagate this effect to the off-diagonal terms in the finite-time transition matrix *T*. For this reason, empirical transition maps alone obscure differences between biases in proliferation and choice towards competing fates, as illustrated in Supplementary Fig. 1**d**.

### Supplementary Note 2: The effect of noisy measurement on transition map inference

In Eq. (5), the transition map is seen to emerge from stochastic state transitions accumulating over time. In practice, an inferred map is also shaped by sources of noise associated with measurement and subsequent dimensionality reduction of the data. In this note, we examine the errors propagated from different technical sources into the observed transition map *T*. As might be expected, we show that technical sources of error lead to a ‘blurred’ transition map, delocalized over the cell state graph. The smoothing kernels connecting the true and observed transition map can be understood as a matrix product of error kernels associated with each individual source of uncertainty.

#### a. Measurement errors

We will consider the errors associated with correctly assigning transition rates from a state *i* at time *t*_1_ to state *j* at time *t*_2_. Such a transition contributes to mass at matrix element *T*_*ij*_(*t*_1_, *t*_2_) of the transition map. At time *t*_2_, errors in measurement re-assign cells from state *j* to another state *k*, with a probability *ϵ*_*jk*_ normalized such that Σ_*k*_ *ϵ*_*jk*_ = 1. With such an error, the observed transition map now becomes 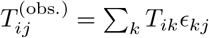. A similar error may occur at *t*_1_. Because technical errors may differ between time points, we will denote *ϵ*^(*i*)^ as the error in measuring the state of a cell at time *t*_*i*_. Accounting for errors in two time points, the observed transition map now becomes:

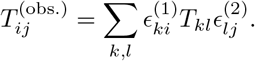

#### b. Clonal dispersion

In LT-scSeq experiments, the cells sampled at *t*_1_ are clonally related to those that give rise to cells sampled at *t*_2_. But being distinct, they may occupy different states. As above, we consider the error in estimating transition rates from state *i* at *t*_1_ to state *j* at *t*_2_. At *t*_1_, a clonally-related state, *k*, is observed instead of state *i*, with a probability that we shall denote *σ*_*ik*_. This probability satisfies normalization Σ_*k*_ *σ*_*ik*_ = 1. Accounting for this clonal dispersion, the observed transition map relates to the true transition map through the relation:

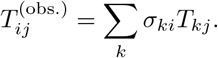

Note that because cells divide, more than one cell may be observed in a clone at time *t*_1_. In this case, the error kernel *σ*_*ki*_ no longer has a unique definition because choices in constructing the transition map may assign more or less weight to particular cells within each clone. By enforcing local coherence, CoSpar strongly weights *σ*_*ki*_ towards states *k* that are close to *i*, thus reducing errors in the transition map as compared to using a ‘naive’ clonal analysis method such as we have previously reported^2^, which weights all cells in a clone at *t*_1_ equally.

Compounding clonal dispersion and measurement error, we recognize the the observed transition map has the form:

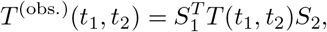

where *S*_1_ = *ϵ*^(1)^*σ* and *S*_2_ = *σ*^(2)^.

### Supplementary Note 3: Coherent sparse optimization

Our goal in dynamic inference is to learn the finite-time transition map, as defined in Eq. (4), for the set of observed cell states in a given experiment. After imposing sparsity and coherence constraints (see main text), we obtain the cost function,

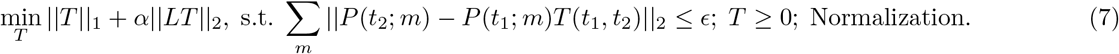

Here, *P*(*t*_1,2_; *m*) is a row-vector representing the distributions of cell states within the *m*-th clone. 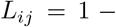 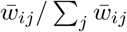 is the normalized graph laplacian, with *w*_*ij*_ the graph connectivity of the nearest neighbor kNN graph of cell states. Defining **P**(*t*) as a clone-by-cell matrix resulting from concatenation of individual clonal distribution: {*P*(*t*; *m*),*m* = 0, 1, 2…}, we note that Σ_*m*_||*P*(*t*_2_; *m*) − *P*(*t*_1_; *m*)*T* (*t*_1_, *t*_2_)||_2_ = ||**P**(*t*_2_) − **P**(*t*_1_)*T* (*t*_1_, *t*_2_)||_2_, which gives the form of the cost function given in the main text. For joint optimization, the cost function is additionally minimized over **P**(*t*_1_), i.e. 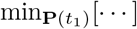.

Before continuing, we note the relationship of this optimization problem to past literature. Absent the coherence constraint (*α* = 0), this optimization problem reduces to sparse optimization by lasso regression. To our knowledge, only one study has explored the extension of lasso to enforce coherence with relation to a data embedding, called ‘fused lasso’ optimization^7^. Fused lasso is however different in three important ways from Eq. (7). First, it suppresses the first-order derivative of the inference target to promote coherence. Second, fused lasso was developed for 1-d or 2-d datasets, assuming a natural ordering for the observed cell states. Third, like lasso, the inference object of fused lasso is a vector. In contrast, the coherent sparse optimization in Eq. (7) is generalized to arbitrary graphs; it suppresses the second-order derivative (the curvature) to enforce coherence; and it is generalized to matrix inference.

Our goal is now to ground the optimization problem in LT-scSeq data, and to propose an algorithm that approximates solution of Eq. (7). To make connection with raw clonal data, we approximate the density profile matrices **P**(t) as,

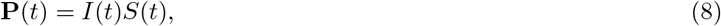

where *I*(*t*) is a clone-by-cell matrix observed at time *t*, and *S*(*t*) is a cell-cell similarity matrix at time *t*. Note that Eq. (8) integrates the state information (encoded in *S*(*t*)) and clonal information (encoded in *I*(*t*)) into **P**. This local smoothing operation indirectly imposes coherent transitions in this system.

We now discuss implementation of the optimization problem. Eq. (7) might be formulated as a quadratic programming problem, and be solved accordingly as in fussed lasso^7^. However, this strategy is very expensive computationally^7^. There could be ways to solve the optimization efficiently and exactly, and we leave it as an open problem. Instead, we provide an efficient yet heuristic way to solve the optimization. Specifically, we break down individual elements of the objective function, and propose a simple alternative for each of them.

#### 1. Sparsification

Instead of including the sparsity term ||T||_1_ into the objective function, we directly apply a pre-defined thresholding to the transition map at each iteration: *T* ← *θ*(*T*, *v*), where

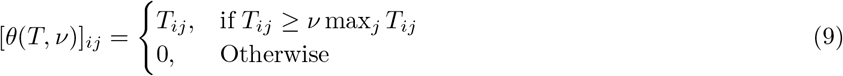

#### 2. Transitions within clones

To enforce Eq. (4) for each observed clone, we consider a clonal transition map π^*m*^ for the *m*-th clone, which allows only intra-clone transitions and conserves the total transition flux within a clone. We do so by projecting the transition map *T* and performing clone-wise normalization: 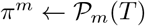:

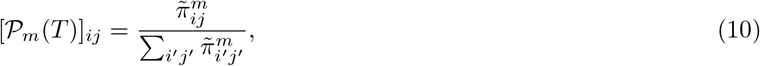

where 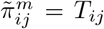 if the transition *i* → *j* occurs within clone *m*, and otherwise 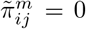. The composite map capturing all intra-clone transitions is then,

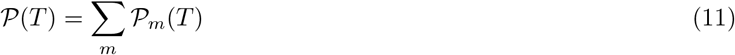

A map constructed in this way, 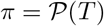, will satisfy the following equation approximately:

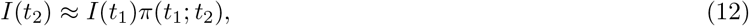

which is the clonal constraint for directly observed cell states^8^. The map *π*(*t*_1_; *t*_2_) can be used to specify *T*, but being constrained to clones it is no longer coherent.

#### 3. Coherence

To enforce coherence, we begin by noting that Eqs. (4), (8) and (12) together lead to the relationship 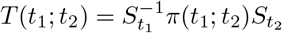. As similarity matrices S are generally non-invertable, we introduce a pseudo-inverse,

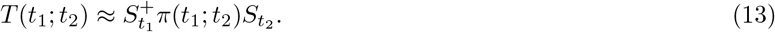

Eq. (13) smoothes the transition map learnt within-clones, *π*, over nearby states to get a transition map *T* across all states. *T* is now a locally continuous map and satisfies the coherence constraint: similar initial cell states have similar fate outcomes.

This approach to calculating *T* leads to minimization of the term *α*||*LT*||_2_ in Eq. (7), although the parameter *α* establishing the relative weight of coherence is no longer explicitly identifiable in the procedure. It is instead reflected in the extent of smoothing.

These three steps, carried out sequentially and iteratively, define the CoSpar algorithm given in methods. Note that normalization is performed clone-wise in Eq. (11). The non-negativity constraint, *T* ≥ 0, is implicitly satisfied in the above steps. In our strategy, Eq. (13) is the most time-consuming step as it involves multiplication of large matrices. CoSpar is nonetheless efficient as it carries out matrix multiplication *only* at Eq. (13), and we find that it converges within a few iterations (Supplementary Fig. 2**d**).

### Supplementary Note 4: Transition map initialization with HighVar

The HighVar method provides an approach to initialize the joint optimization of *T* and *I*(*t*_1_) (see Methods). The approach is loosely motivated by the expectation that cells similar in gene expression between time points may share clonal origin. This expectation can be violated; we use it only to initialize numerical optimization.

HighVar consists of three steps: 1) Select highly variable genes that are expressed at both *t*_1_ and *t*_2_; 2) For each highly variable gene (indexed by *m*), threshold its expression to form a binary expression matrix 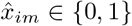 for all states observed at *t*_1_ and *t*_2_ to generate pseudo clonal data 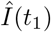 and 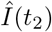 from the binary expression matrix; 3) Run CoSpar with 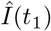 and 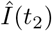. The pseudo-clonal data 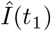 and 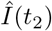 are discarded, and the resulting map *T* is used to initialize CoSpar with the true clonal data.

For the first step, we use the SPRING gene filtering function filter genes with an adjustable gene variability percentile parameter HighVar gene pctl to select highly variable genes^9^. For the second step we discretize the gene expression of each highly-variable gene, sequentially, with a gene-specific threshold *η*_*m*_:

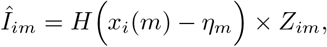

where *H*(·) is the Heaviside step function (*H*(*x*) = 1 if *x* > 0; otherwise 0), 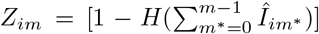 sums over previously considered genes to ensure that the same cell is not assigned to more than one pseudo-clone. The gene-specific threshold *η*_*m*_ is chosen such that every pseudo clone has the same number of cells at each time point *N*_*t*_/*M*, where *N*_*t*_ is the number of observed cells at time *t* and *M* is the total number of highly variable genes (i.e., pseudo clones). In case *N*_*t*_/*M* is not an integer, we use its ceil, i.e., [*N*_*t*_/*M*], and stop the clonal matrix update when all cells are clonally labeled.

